# Are online corrections to visual targets really a distinct class of movement?

**DOI:** 10.1101/2025.03.14.643316

**Authors:** David Y. Mekhaiel, Melvyn A. Goodale, Brian D. Corneil

## Abstract

Humans have a remarkable capacity to adjust reaching movements rapidly and accurately when visual targets jump to a new location. The short latency of such online corrections has led to the hypothesis that they constitute a distinct class of movement and arise from an ‘automatic pilot’ that is selectively engaged only during ongoing movements.

Here, we test this idea by measuring muscle recruitment, force, and kinematics in a jumping target reaching task. In separate blocks of trials, participants were instructed to respond to target jumps by (1) following the jumped target, (2) stopping the on-going movement, or (3) ignoring the jumped target. This allowed us to establish the automaticity and timing of responses to target jumps and to compare such measures to the original reaching movement initiated from rest.

We find that the earliest phase of muscle recruitment elicited by the jumped target corresponds to a subcortical reflex, beginning at ∼80ms and ending by ∼120ms, preceding the onset of voluntary recruitment at ∼130ms. This reflex inexorably drives a reaching adjustment towards the new target in all three blocks; it is only somewhat reduced in the ‘stop’ and ‘ignore’ blocks. Critically, this earliest phase of muscle recruitment was also present at the exact same latency (80ms) for the original reaching movement initiated from rest. Thus, rather than supporting the model of online corrections as distinct class of movement that is mediated by an ‘automatic pilot,’ our results suggest that all reaches, whether adjusted in mid-flight or initiated from rest, arise from a common nested control system featuring subcortical and cortical components whose influence can be strategically preset by task demands. Our results also reinforce the importance of considering movement biomechanics when interpreting kinematic latency differences across movements made in different situations.

## Introduction

It has long been recognized that adjustments in movements in response to suddenly displaced targets, often termed ‘online corrections,’ can be initiated at shorter latencies than movements initiated from rest (see Smeets and colleagues 2016^1^ and associated commentaries). While the reported reaction times for these types of movements can be quite variable depending on stimulus properties, task instruction, and methods of analysis, the short-latency and inexorable stimulus-directed nature of the initial phases of online corrections are frequently noted as hallmarks distinguishing online corrections from other types of movements (for review, see Gomi^2^). These differences are often attributed to a rapid ‘automatic pilot,’ which exclusively modifies movement plans mid-flight but is not engaged during movement from rest^3^. While online corrections undoubtedly have unique kinematic features, debate persists regarding the neural correlates of online corrections, with theories suggesting that processing through either the posterior parietal cortex (PPC) ^3,4^ or the superior colliculus (SC) ^5,6^ instantiates the ‘automatic pilot.’ Regardless of the exact neural underpinnings, it seems curious that the brain would have a dedicated system that is utilized only in the case of online corrections; if such a rapid system exists, why is it not used for movements from rest? And, what determines the stimulus-driven nature of the first phase of online corrections ^7,8^?

We designed an experiment to study and compare online corrections and movements from rest, using electromyography (EMG) to characterize the timing and pattern of muscle recruitment that precede both types of movement. We also measured reach force using force transducers in the reach manipulandum. Although early investigations of online corrections used EMG measures to help determine the latency of when an online correction is commanded^9^, this has been more the exception than the rule. Yet, the complementary use of EMG and measures of force can also help clarify the impact of movement biomechanics, which may well be quite different for a movement initiated from rest versus an online correction, due for example to recruitment changes on actively lengthening muscles. Thus, in the present experiment, across three separate blocks of trials, participants performed different reach adjustments in response to a target that jumped unpredictably on a third of all trials to a second location. Importantly, all blocks featured identical visual stimuli, ensuring that observed differences in EMG, forces, and movement kinematics stemmed solely from the task set.

In one block, participants were instructed to ‘Follow’ the target jump by performing an online correction. This allowed us to compare the timing of muscle recruitment leading up to movements initiated from rest (during the initial reach) and online corrections (during the reach adjustment); an ‘automatic pilot’ predicts earlier recruitment during online corrections. In another block, participants were instructed to ‘Stop’ their ongoing movement and hold their hand in place if the target jumped. Although some studies suggest that muscle recruitment for action-stopping occurs later (∼140ms)^10–12^ than for online corrections, other studies have suggested that action-stopping may be produced by the same rapid neural network^13,14^. If so, the first phase of muscle recruitment in the ‘Follow’ and ‘Stop’ blocks should start at the same time, but be linked to the different goals of these tasks. In another block, participants were instructed to ‘Ignore’ the target jump and continue their reach towards the original target location. This block allowed us to test the automaticity of the muscle activity underlying online corrections.

Our results indicate that the earliest phase of muscle recruitment elicited by the target jump arises from a subcortical reflex that produces express visuomotor responses (EVR; formerly termed ‘visual’ or ‘stimulus-locked’ responses^15,16^; see Contemori et al.^17^ for rationale behind the “express” terminology) spanning an interval from 80-120ms after target jump, prior to the onset of voluntary recruitment at ∼130ms. Upon comparing the reaches initiated from rest and the online corrections, we found that the timing of both the EVR and forces applied to the manipulandum were virtually identical. Online corrections, however, had much lower kinematic reaction times than movements initiated from rest, likely because biomechanical considerations delayed how quickly consequent forces overcame the arm’s inertia during movements from rest compared to online corrections. Remarkably, the timing and patterning of the EVR did not change with task set. EVRs inexorably drove the online correction towards the jumped target in all three blocks; it was only somewhat reduced in the ‘Stop’ and ‘Ignore’ blocks. EVR magnitude was modestly (r^2^ = ∼.25) correlated with reaction time across all blocks, eliciting short-latency force and kinematic online corrections towards the target appropriately in the ‘Follow’ block and erroneously in the other two blocks. The similarity of the EVR across all blocks suggests that it is the likely neuromuscular correlate that initiates online corrections, explaining why the reaching arm is inexorably drawn initially towards displaced targets ^8,18^. Overall, rather than supporting the idea of an ‘automatic pilot’ that is exclusively engaged during online corrections, our results suggest that the unique kinematic features of online corrections can arise from a common nested control system featuring subcortical (EVR) and cortical (voluntary muscle activity) components that, depending on behavioral context, can be similarly active during both online corrections and reaches from rest.

## Results

### Overview

During intermixed blocks of trials, participants were instructed to adjust their ongoing reaches differently in response to the exact same visual stimuli by performing three types of reach adjustments. Targets initially appeared randomly in one of two locations (T1 or T2). Participants were instructed to reach towards this initial target as soon as it appeared. On a third of trials, the target jumped to the other location once the participant’s hand left the start point (fig. 1C; within 8ms of the hand leaving the target). When the target jumped during ‘Follow’ blocks (fig. 1E), participants were instructed to reach towards the NEW target; when it jumped during ‘Stop’ blocks (fig. 1F), they were instructed to stop the ongoing movement and hold their hand in place; and when it jumped during ‘Ignore’ blocks (fig. 1G), they were instructed to continue reaching towards the OLD target position. No explicit feedback was provided about what the duration of the movement should be.

**Figure 1.**
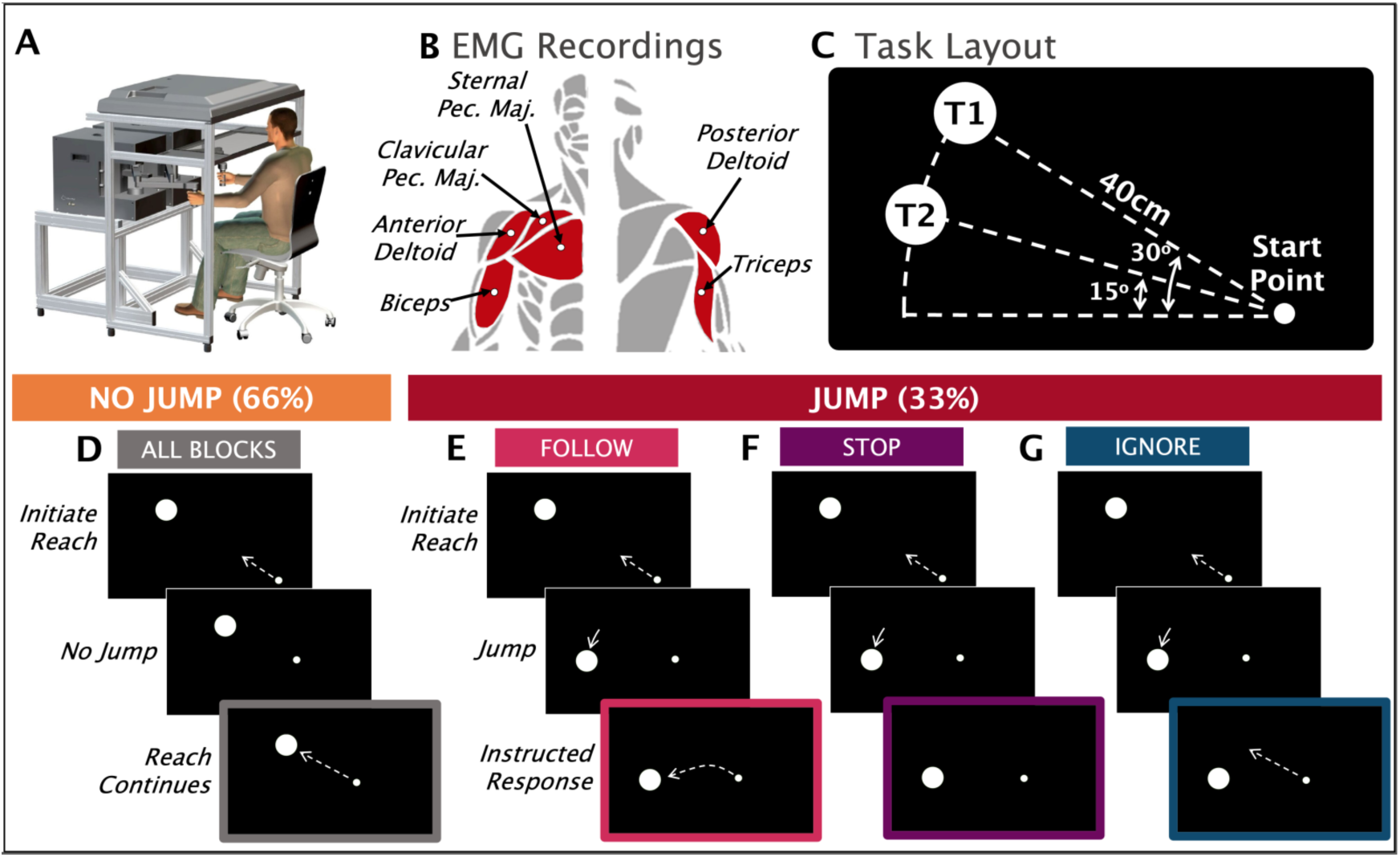
Experimental apparatus & task: (A) Participants made reaches with their right arm in a Kinarm End-Point Lab. (B) Surface EMG activity was recorded from six right arm muscles. (C) The task layout included a startpoint and two potential targets (T1 and T2). (D-G) Task timelines. Solid arrows denote target movement and dashed arrows denote hand movement.

Below, we compare the latency of the mean kinematics, force, and EVR onsets during the initial reach (in response to the initial target) and during the online correction (in response to the target jump), report kinematic and mean force measures obtained from the Kinarm manipulandum (fig. 1A) and describe how error rate and reaction time of force deviation varied across different blocks. We then report surface EMG signals recorded from arm and shoulder muscles (fig. 1B) to investigate how the task set influences the EVR and how the EVR relates to reaction time of force deviation.

### Similar force onset for online corrections and reaches from rest

We focus first on comparing kinematic reaction times and the timing of changes in manipulandum force during the initial reach compared with those observed in the online reach correction. Past work reporting shorter latency kinematic reactions times for online corrections employed either visual inspection ^5,19–21^ or various acceleration and velocity thresholds ^22–26^. Consistent with previous findings, if we apply a common threshold of 2m/s^2^ for acceleration onset, we observe longer kinematic reaction times for the initial reach of 202ms (to T1) and 195ms (to T1), compared to earlier kinematic reaction times for online correction of 147ms (T1 to T2) and 145ms (T2 to T1; fig. 2). Clearly from fig. 2, at least a portion of this difference appears to reflect how rapidly the acceleration traces rise to this threshold. Even so, from visual inspection, it does appear that acceleration rises earlier during online corrections (∼100ms) than it does for reaches from rest (∼150ms); for alternatives for the determination of onsets that do not rely on arbitrary thresholds, see Wijdenes et al.^27^.

**Figure 2.**
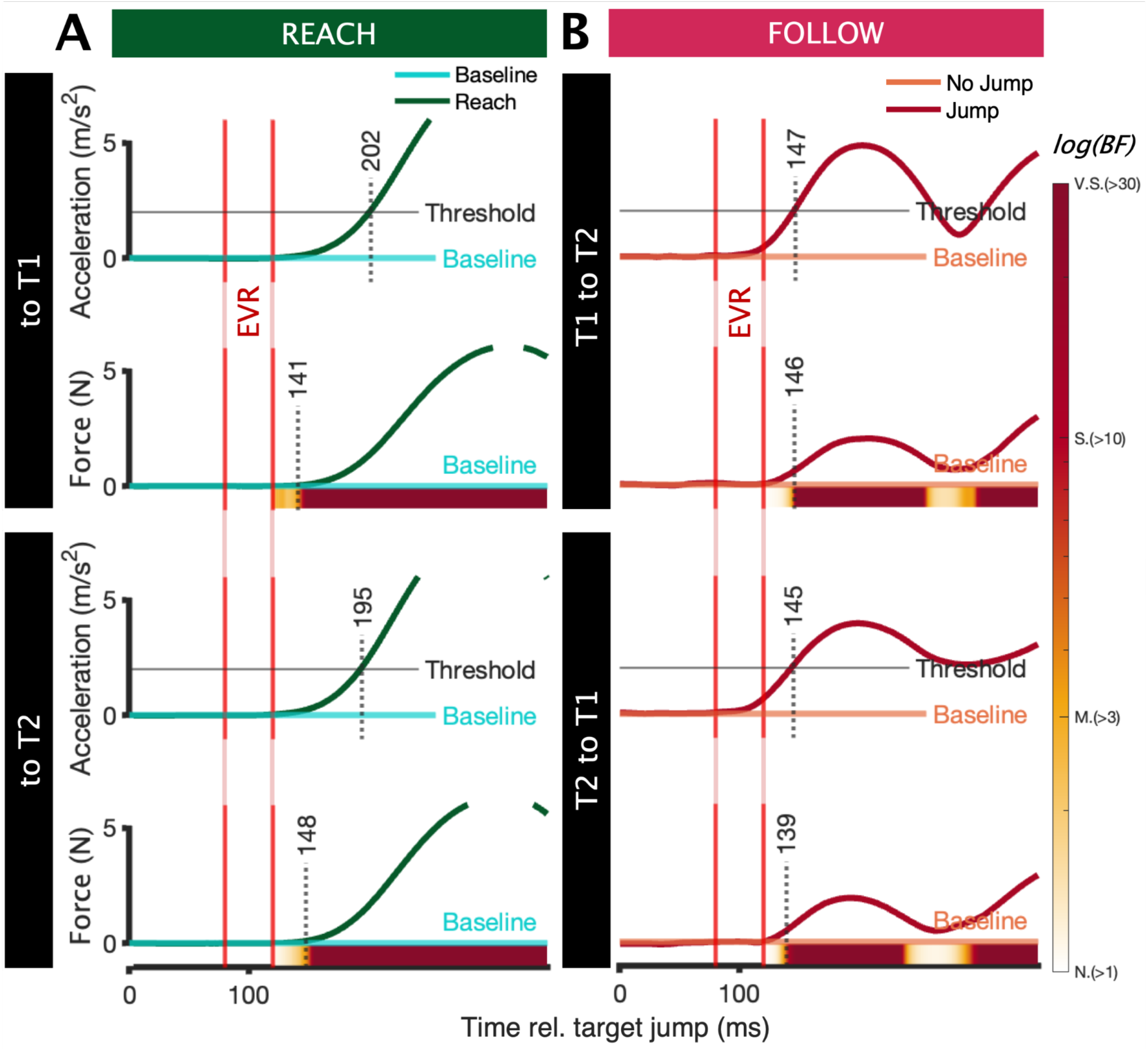
Comparison of reaction times determined from movement kinematics vs kinetics: Mean acceleration (vector sum of x and y acceleration) and mean force deviation from baseline for (A) initial reach and (B) online reach correction. For acceleration plots, dotted vertical lines denote onset determined using a threshold of 2 m/s^2^. For force plots, the Bayes’ Factor (BF) heatmaps denote the likelihood of the model that the force deviates significantly from baseline (for the initial reach comparison begins at 120ms, force 200ms prior was used as baseline). Dotted vertical line denotes first instance where BF > 10. BF heatmap is qualified by the following standard thresholds ^30^: N.(> 1) = Negligible, M.(> 3) = Moderate, S.(> 10) = Strong, V.S.(> 30) = Very Strong. EVR epoch (80-120ms) denoted by vertical red lines.

We followed up with a second analysis, in which we employed a more sensitive timewise Bayesian analysis to determine force onset in both the initial movement and the correction (fig. 2). This analysis compares the likelihood of divergence between the mean force recruitment during the initial reach and online correction at each millisecond, and follows suggestions that measures of force may provide a better method for measuring reaction times^28^. Additionally, the Bayesian analysis does not rely on an arbitrary threshold, relying instead on the variability of movements from earlier points in time. Furthermore, this analysis is non-parametric and arguably does not require false discovery rate corrections in the traditional sense required for frequentist approaches^29^. Finally, we denoted onset as the first instance of sustained divergence using a conservative threshold of BF>10, indication a strong likelihood of a difference.

To determine the latency of online reach corrections, we compared force data on jump vs no-jump trials. To determine the latency of the reach from rest, we compared force of the initial reach to baseline force (200ms prior). Using this analysis, we found that the latencies for force were nearly identical across both types of movements; 141ms (to T1) and 148ms (to T2) for reaching from rest compared to 146ms (T1 to T2) and 139ms (T2 to T1) for online correction (fig. 2). This analysis reinforces previous findings that the method used to determine response latency can have a large impact on the conclusions that may be drawn from such analyses. Although more traditional methods of identifying latency found that online corrections occur earlier than reaches from rest, changes in the mean force applied to the manipulandum begin nearly simultaneously for online correction vs movements initiated from rest.

### Similar EVR latencies for online correction and reaching from rest

The previous analysis suggests that the timing of changes in mean force applied to the manipulandum are not different for online corrections vs movement from rest. It is possible, however, that this result could arise from the subset of faster responding trials and/or subjects. Thus, as a next step we looked at within trial changes in muscle recruitment (Fig. 3), constructing ‘heat maps’ for an exemplar participant that depicts the magnitude of muscle activity on each trial relative to the moment the hand left the start point in response to the initial target appearance (left two subplots, Fig. 3A) or relative to the timing of the jump in the ‘Follow’ block (right two subplots, Fig. 3A). When represented in this way, a distinct band of muscle activity during the EVR epoch time (80-120ms) becomes obvious, consisting of a stimulus-locked increase (for movements from rest, or for when T1 jumps to T2) or decrease (when T2 jumps to T1) in muscle activity. This is the EVR ^16^, as its timing is more aligned to stimulus onset rather than movement onset, and since it facilitates arm movement towards the new target. We find that as with EVRs during online corrections, the initial reach EVRs consistently begin at a latency of ∼80ms, and in both cases the EVR is more aligned in time to stimulus onset/jump rather than F-RT. Although the EVR appears to be larger for online corrections, we note the substantial differences in EMG activity immediately before the EVR. Regardless, a key prediction of the ‘automatic pilot’ hypothesis is that EVR and force changes should be earlier for online reach corrections versus reaches from rest. Although one study has previously reported this when comparing data collected across two different blocks of trials ^31^, another study has also documented nearly identical latencies for EMG onset for both the initial reach and the online correction ^9^. Importantly, our paradigm is designed to permit a direct comparison of EVR timing within the same trials, thereby controlling for target relative to hand location as well as task set.

**Figure 3.**
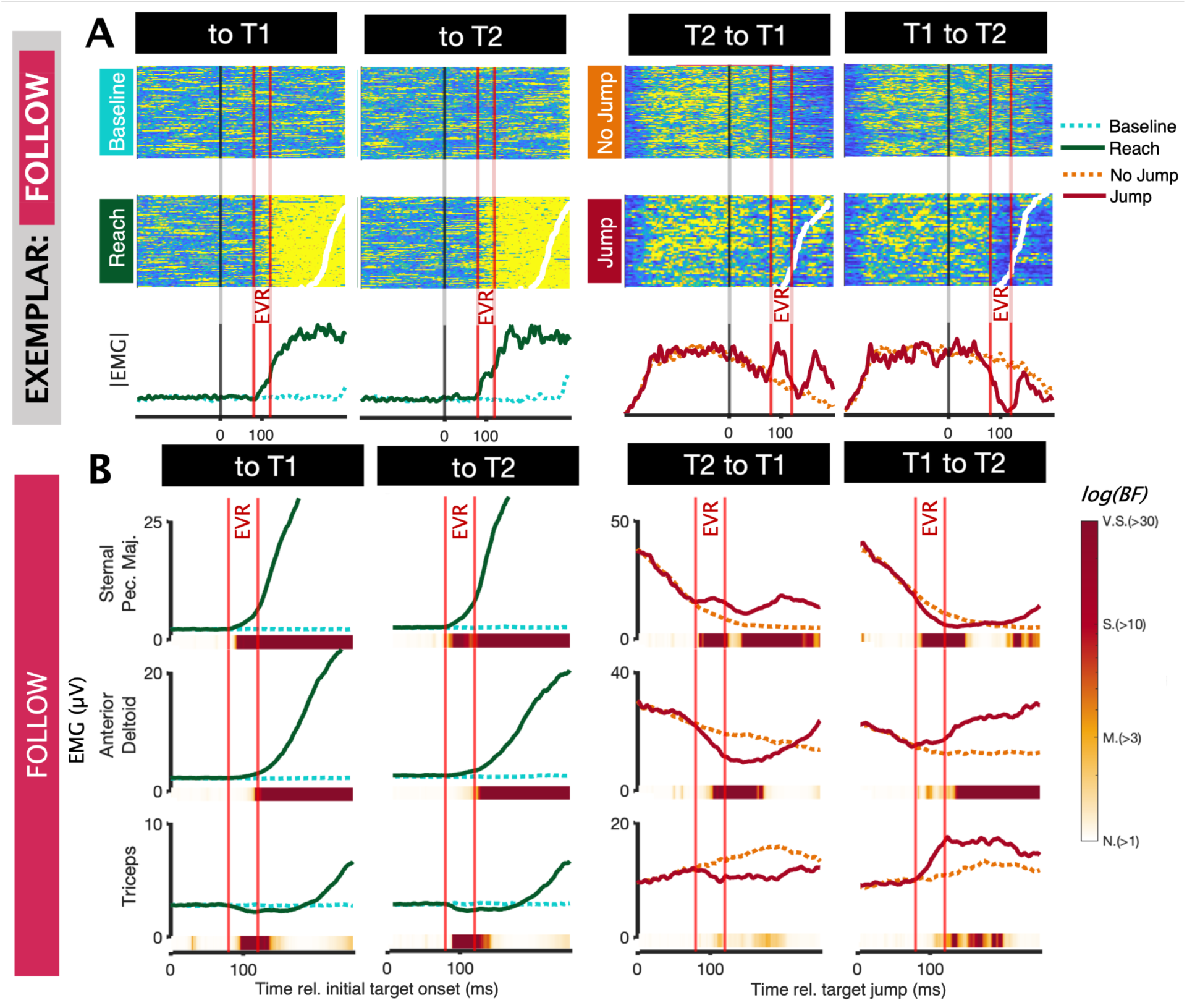
Comparison of latency of change in EMG activity during the initial reach and online correction: (A) Initial reach and baseline single trial heatmaps and means of normalized EMG of the right sternal head of pectoralis major of the exemplar participant (p6). Trials are organized by F-RT, as denoted by the white dots. (B) Group initial reach compared to baseline (200ms prior to target onset) and jump compared to baseline (no jump trials) in ‘Follow’ blocks. Note differences in y-axis scaling. BF heatmap denotes the likelihood of a difference between two conditions. Standard BF thresholds ^30^: N.(> 1) = Negligible, M.(> 3) = Moderate, S.(> 10) = Strong, V.S.(> 30) = Very Strong. EVR epoch (80-120ms) is denoted by vertical red lines.

To compare the latency of the EVRs, we ran a paired Bayesian t-test on each time point, comparing mean EMG in jump vs. no jump conditions and Initial reach vs. baseline (time point 200ms prior; fig. 3B). We found that EVR latency was essentially the same for both the initial reach and online correction. For the sternal head of pectoralis major, the primary muscle used to make right arm leftward reaches, the BF indicated a strong likelihood of an effect starting on average (across jump directions) at 87ms and 85ms during the initial reach and online correction, respectively. Results were less clear for the anterior deltoid and triceps, given that recruitment of these muscles is relatively modest for movements from rest toward either T1 or T2. However, there do seem to be small increases (for anterior deltoid) or decreases (for triceps) during the EVR interval for movements from rest, resembling in time the larger changes in these muscles during online corrections. Overall, our EMG results illustrated in fig. 3 add to our analyses of manipulandum force illustrated in fig. 2 by showing the trial-by-trial consistency in the arrival of motor commands at the muscles, with a latency of ∼85 ms for both reaches from rest and online corrections.

### Analysis of performance

Next, we turn to a comparison of the EVR to the jumped target across the different ‘Follow’, ‘Ignore’, and ‘Stop’ blocks. We first quantified the error rate and reach trajectory to ensure participants followed block instructions. To clarify, error trials were *not excluded* from future analyses. Given the different instructions in each of the three block types, the criteria for determining an error differed for each block (for exemplar, see fig. 4A). Error trials were defined as the participant reaching the original target in ‘Follow’ blocks, reaching either target during ‘Stop’ blocks, or reaching the new target in ‘Ignore’ blocks. Success rates, although defined differently, were highest in the ‘Ignore’ blocks (94.8%), followed by the ‘Follow’ (86.8%), and ‘Stop’ blocks (66.4%; fig. 4C). This largely indicates that participants were attempting to perform the task as instructed. It also suggests that countermanding during the ‘Stop’ condition was the least automatic of the three reach adjustments. Errors were slightly higher when the target jumped away from the participants (T2 to T1; 21.5%) compared to when it jumped towards the participant (T1 to T2, 13.1%, fig. 4B), perhaps reflecting a marginally more difficult adjustment (fig. 4B).

**Figure 4.**
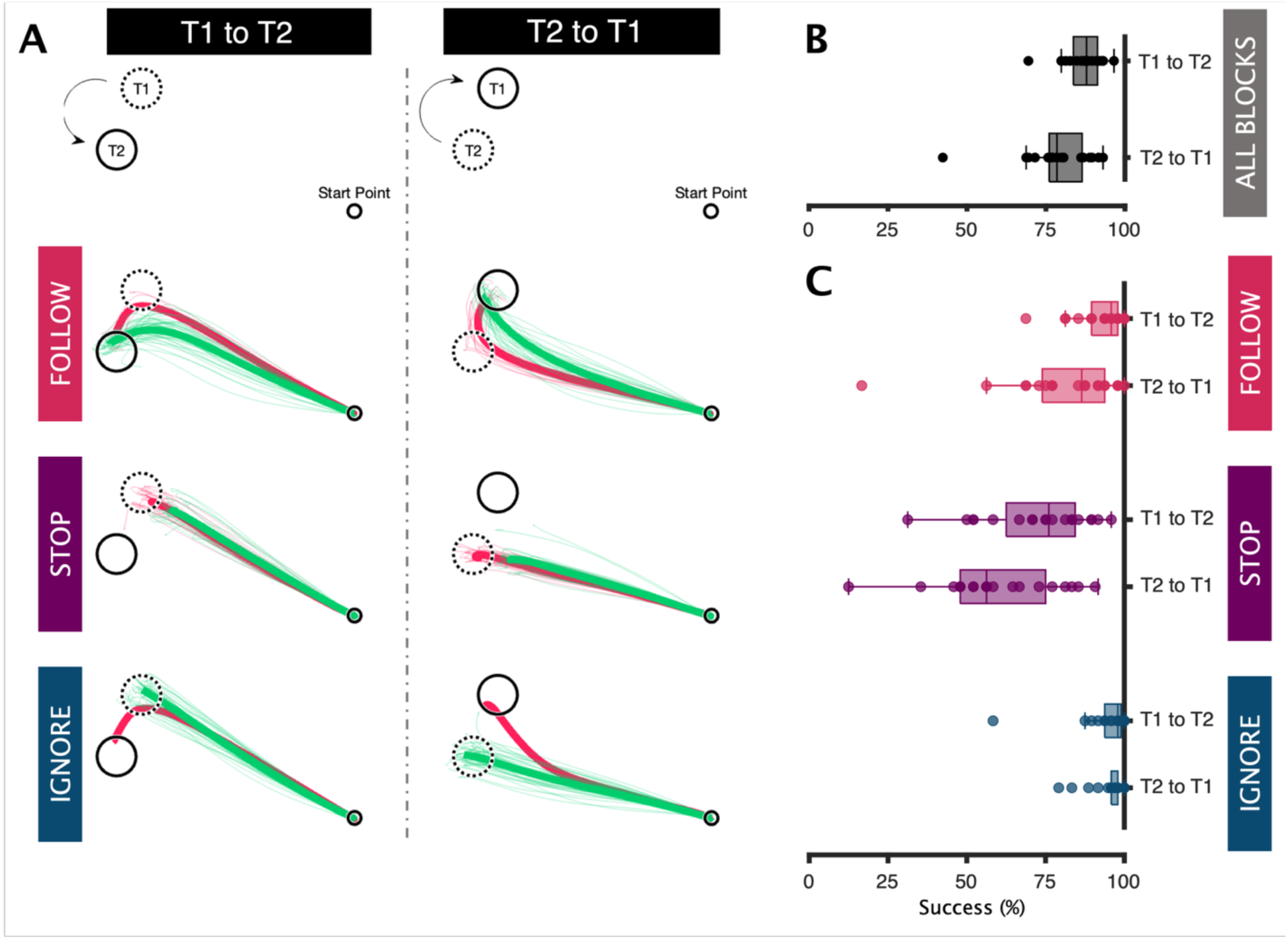
Reach performance across jump trials in different blocks. (A) Reach trajectories for correct (green) and error (red) jump trials for the exemplar participant (p6). Success rate by (B) direction of jump and (C) block instruction. Success rate was determined using criteria unique to each block type, reflecting the unique demands for how participants should respond to the jumped target.

### Initial online correction observed regardless of instruction

We then looked at force and reaction times to compare responses across blocks. To make interpretation of the force and kinematic data easier, we rotated the workspace by −157.5° around the center of the startpoint (fig. 5A). Doing so aligns the main force moving the arm towards the average position of the two targets with the x-axis and the direction perpendicular to the average direction of the two targets relative to the startpoint with the y-axis. Looking first at the force trajectories during the ‘Follow’ block, we find that the force trajectories appear to diverge in the y-dimension near the end of the EVR epoch (∼110ms; fig. 5B). At this point, there does not appear to be force deviation in the x-dimension, meaning participants did not notably slow down their outbound movement following a target jump. We observed similar but smaller force deviations in the y-dimension both in the ‘Stop’ and ‘Ignore’ blocks, emphasizing a seemingly reflexive deviation in reach force toward the jumped target. In the ‘Stop’ block only, was there also a clear force deviation in the x-dimension starting at a latency of ∼150ms, ∼40ms later than the onset of online reach corrections (∼110ms). Therefore, in the ‘Stop’ block, participants stopped their ongoing reaching movement only after an initial deviation toward the jumped target.

**Figure 5.**
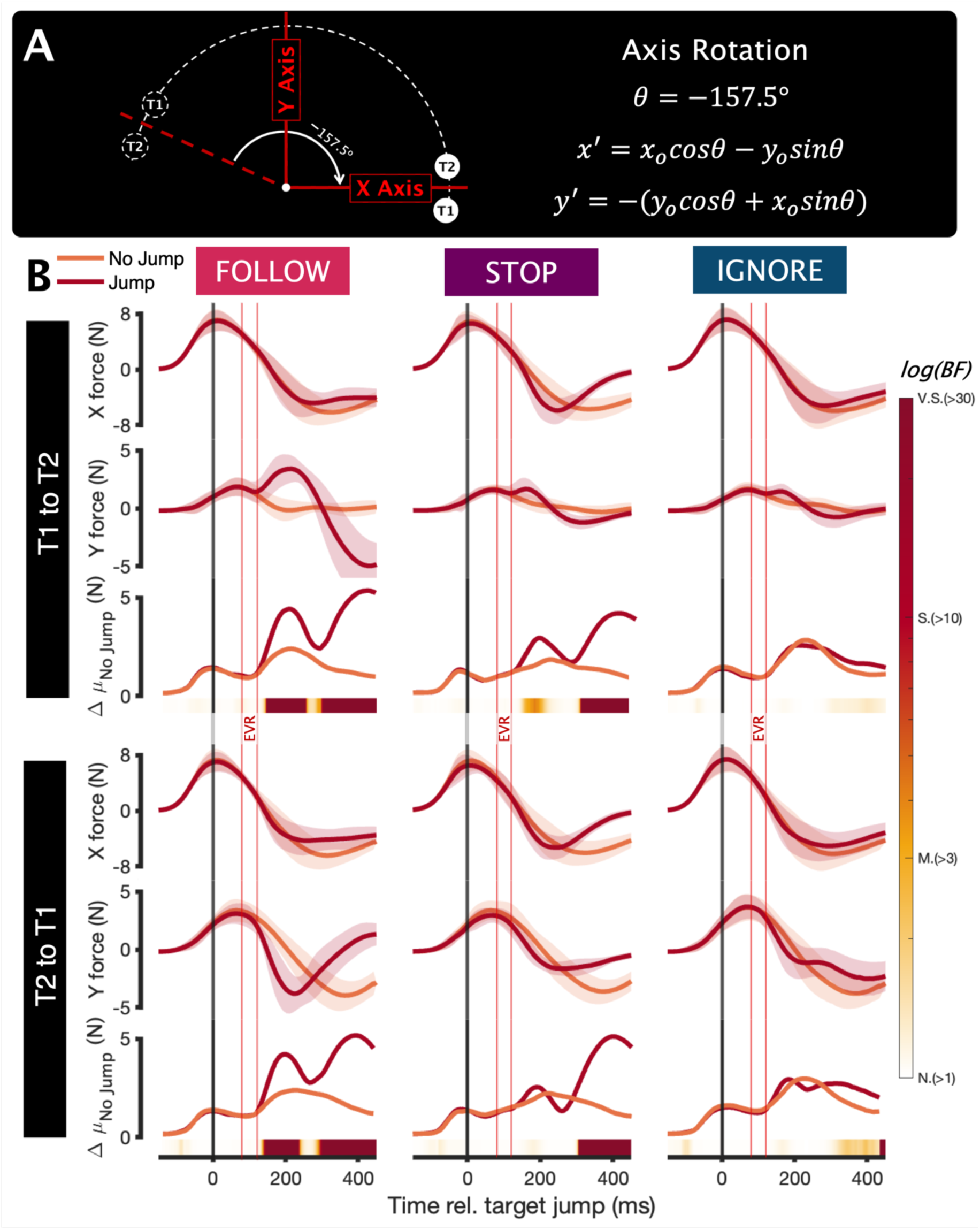
Evolution of force applied to manipulandum through time. (A) Alignment of force data to isolate movement components between startpoint and targets (x-dimension) and deviation towards each of the two targets (y-dimension). (B) Mean group force trajectories, mean force deviation, and Bayesian analysis. For X and Y force trajectories, boundaries indicate standard deviation across participants. To obtain Δµ_No Jump_, we compared each participant’s trial-by-trial trajectories to their mean no-jump trajectory, across both jump and no-jump trials. The resulting standard deviations were then averaged across participants. BF heatmap denotes the likelihood of the model that jump trials exhibit more deviation from the mean no jump force trajectory, than no jump trials. BF heatmap is qualified by the following standard threshold, as per the legend ^30^: N.(> 1) = Negligible, M.(> 3) = Moderate, S.(> 10) = Strong, V.S.(> 30) = Very Strong. EVR epoch (80-120ms) is denoted by vertical red lines to demonstrate the onset of EVR responses (first line) and the onset of voluntary muscle activity (second line).

To better characterize these observed differences, we then examined how the target jump impacted force development over time. We used a Bayesian paired t-test to compare the mean deviation from control (mean no jump) for both jump and no jump conditions for each time point. In ‘Follow’ blocks, for each jump direction, there begins to be a ‘strong’ likelihood of divergence (*BF* > 10) at 140ms (T1 to T2) and 134ms (T2 to T1) and in ‘Stop’ blocks at 151ms (T1 to T2) and 303ms (T2 to T1; a small divergence at ∼150ms approached but did not reach statistical threshold). In ‘Ignore’ blocks, trajectories strongly diverged in only one jump direction (T2 to T1), starting at 321ms. These results demonstrate that our task elicited significant short latency corrections in both jump directions in both the ‘Follow’ and ‘Stop’ blocks. Furthermore, countermanding or action stopping appears to begin later than online correction, at a similar latency to voluntary reaching from rest.

To better characterize the aforementioned small deviations in the force trajectories, we next investigated how force reaction times (F-RT) differed across conditions. F-RT denoted the first time point in each trial where the force trajectory was more than one standard deviation away from the mean force trajectory of the complimentary no-jump condition. Therefore, here F-RT is defined agnostically, rather than specifically reflecting block goals. For example, in ‘Stop’ blocks a F-RT could indicate either a correct stopping action or an erroneous online correction. Since this method of reaction time detection is likely prone to false positives, especially during an ongoing movement, we also conducted the same analysis for no jump trials to serve as control.

Comparing the distributions of F-RTs in jump and no jump trials, we found that we began observing more ‘F-RTs’ in jump conditions relative to no jump conditions starting at ∼100ms after target jumps during ‘Follow’ blocks (fig. 6). Looking at how those reactions impacted movement, we found that for all blocks the earliest F-RTs (100-150ms) typically resulted in acceleration towards the jumped target (below the control line). This was most prominent in the ‘Follow’ and ‘Stop’ blocks, and to a lesser degree in the T2 to T1 jumps during ‘Ignore’ blocks. Although this is expected in ‘Follow’ blocks, this is counter to the task goals in both the ‘Stop’ and ‘Ignore’ blocks. This observation reiterates that online corrections (at least within our task) seem to have a reflexive component that cannot be arbitrarily suppressed by task set.

**Figure 6.**
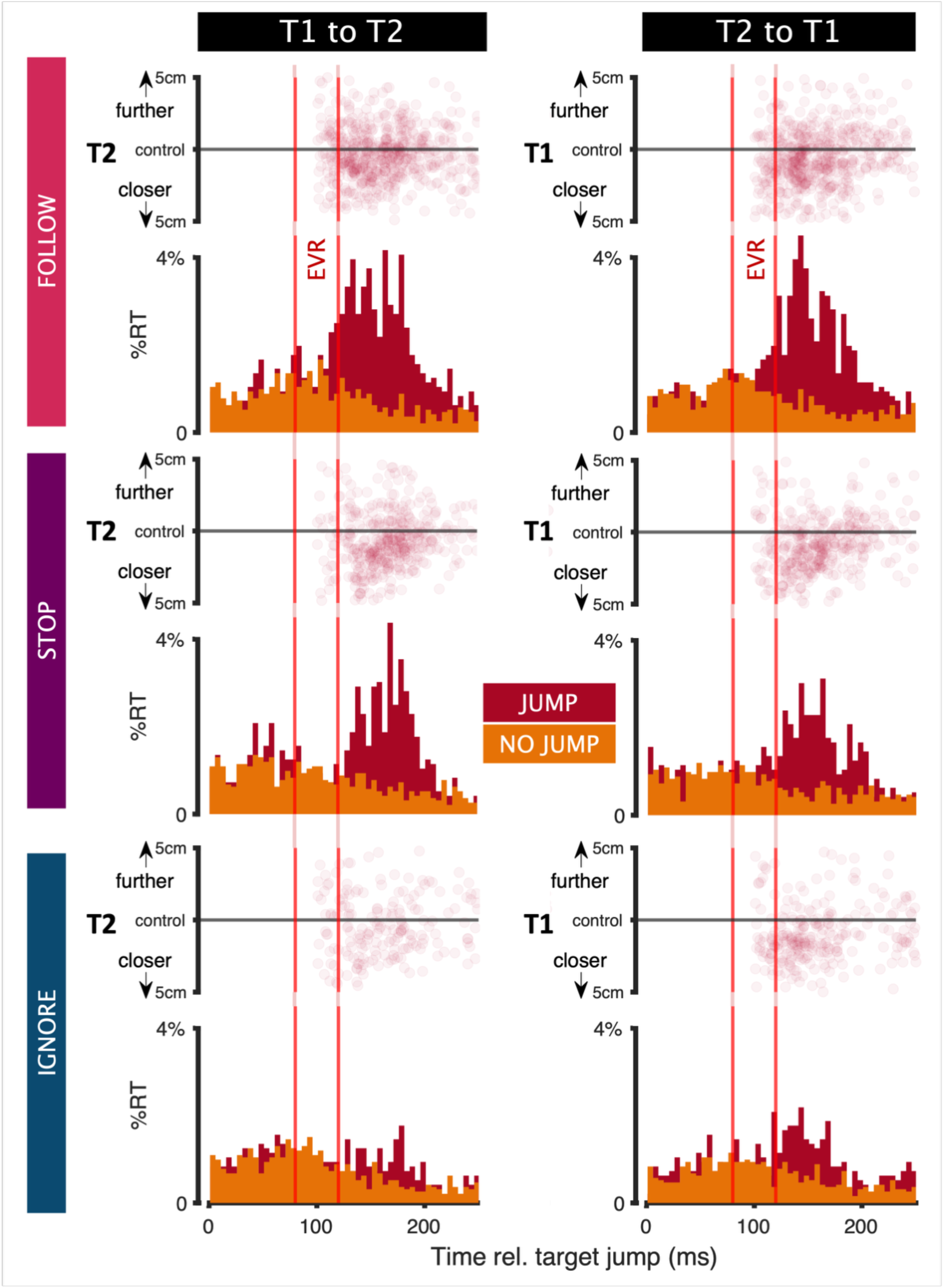
Distribution and direction of F-RTs: Each of six subplots displays histograms of all F-RTs for all participants (5ms bins) for both jump and no jump conditions as a percentage of trial type. F-RTs determined as the first instance where the force trajectory deviates from the no-jump force trajectory by more than one standard deviation for at least 1ms. Scatter plots denote the deviation on an individual trial basis relative to each of the two targets (T1 & T2) 20ms following F-RT, minus the distance to each target for the corresponding mean no jump condition. A dot above the control line denotes movement towards the target relative to the control condition and above the line denotes movement away from the target.

### EVR present in all three tasks

Next we examined single-trial EMG recordings, to better understand the changes in muscle recruitment related to the profiles of reach forces. Again, we represent EMG activity as heat maps, doing so first for an exemplar participant (fig. 7). This revealed that, in all blocks, the jumped target elicited prominent EVRs, with obvious stimulus-locked increases (when T1 jumps to T2) or decreases (when T2 jumps to T1) in pectoralis muscle activity. In the exemplar participant, these directional changes in muscle activity begin ∼80ms after the target jump. Like online motor corrections ^8^, EVRs are known to be spatially locked to stimulus location independent of task ^32,33^. The consistency of the EVR across blocks strongly implies that the initial online corrections observed across blocks (fig. 6), are attributable to the EVR, since it is the only distinct phase of muscle activity occurring prior to those short-latency corrections. Importantly, while the magnitude of the EVR seems to be somewhat muted in the ‘Ignore’ block for this participant, it clearly persists even on a trial-by-trial basis.

**Figure 7.**
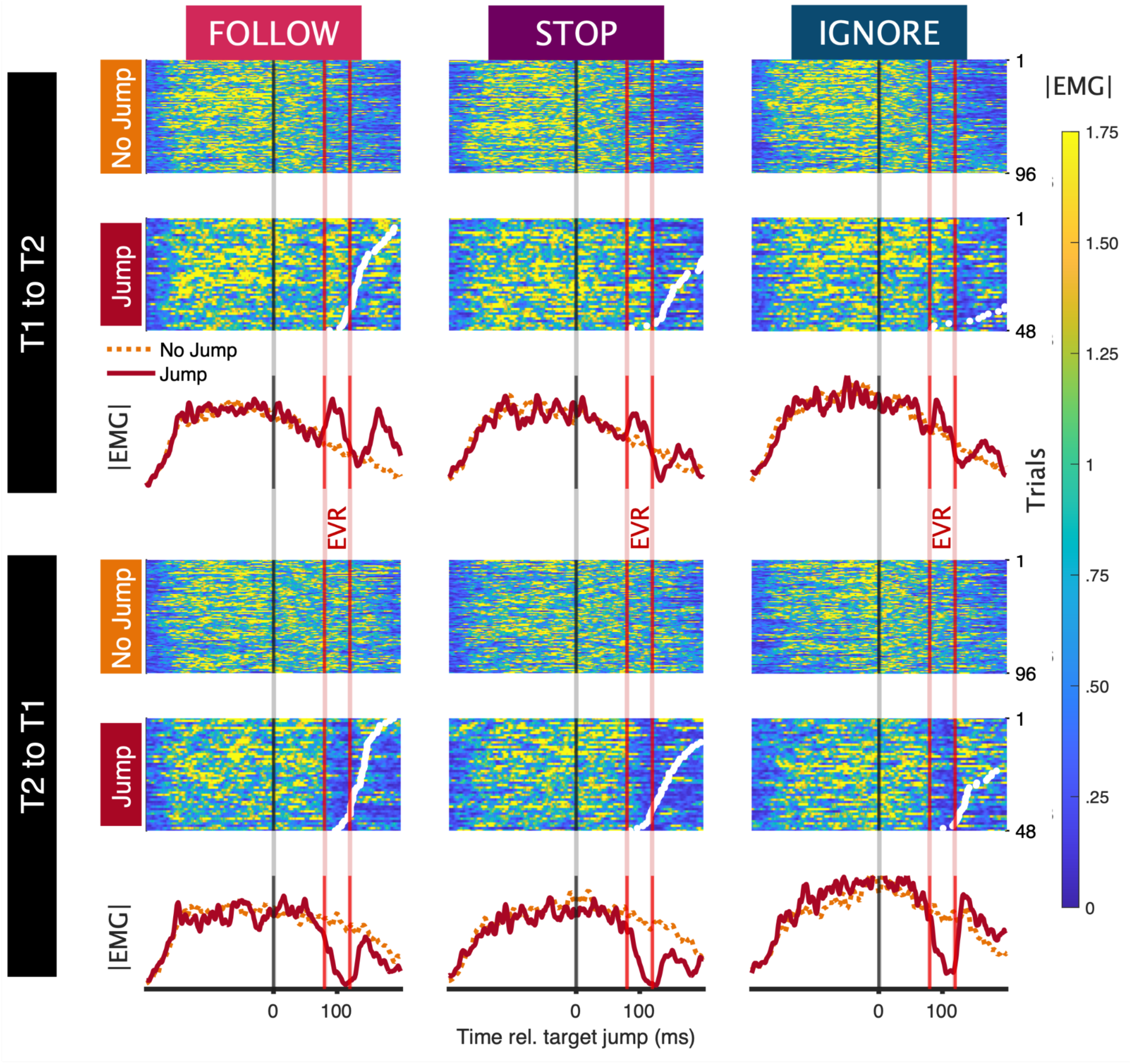
Representative example of EMG activity recorded in jump vs no jump trials in all blocks: Each of six subplots show single trial heatmaps and means of normalized EMG of the right sternal head of pectoralis major of the exemplar participant (p6), aligned to target jump (or, for no jump trials, when the target would have jumped). Jump trials are ordered by F-RT, as denoted by the white dots. EVR epoch (80-120ms) is denoted by vertical red lines.

### Instruction modulates EVR magnitude but not timing

Next, we sought to determine if the magnitude of the EVR differs across block types. As seen in the mean EMG plots (fig. 8A), muscle activity for no jump and jump conditions is apparent across all muscles starting at ∼80ms. Notably, the EVR magnitude typically changes in the same direction as the subsequent muscle activity during the ‘Follow’ blocks, therefore facilitating arm movement in the direction of the new target. This means that across the two jump directions (T1 to T2 vs. T2 to T1), we observe an inversion of the direction of the EVR relative to the control no jump condition.

**Figure 8.**
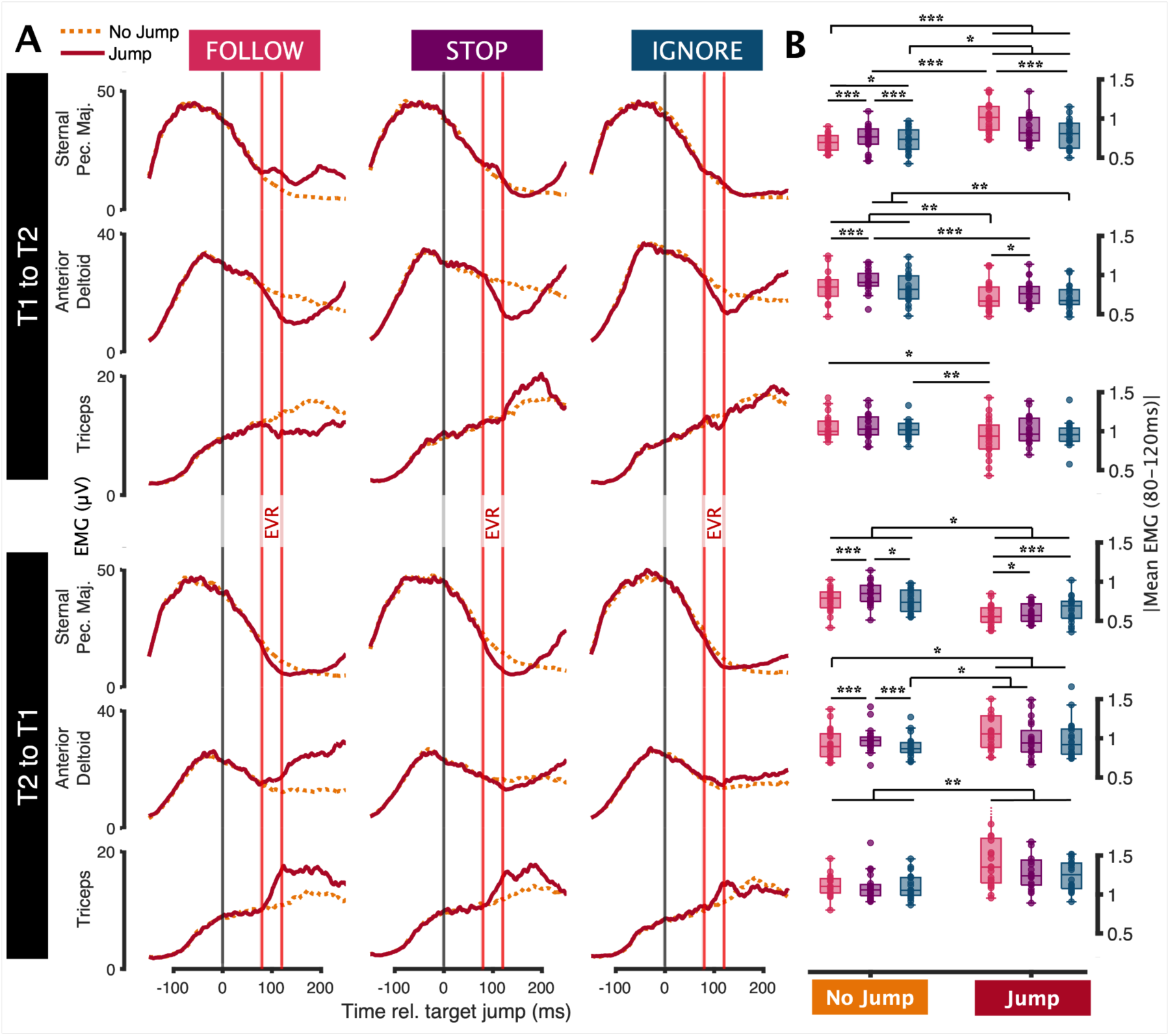
Representation of EMG activity across the sample, for different muscles and blocks: (A) Mean EMG of all participants across three muscles, for both no jump and jump conditions, including error trials. EVR epoch (80-120ms) is denoted by vertical red lines. (B) Mean EMG magnitude during the EVR epoch. Asterisks indicate significant findings using a contrast of estimated marginal means based on a linear mixed effects model. Due to the size of the dataset, degrees of freedom were estimated asymptotically. The Tukey method was used to adjust p-values. Grouped significance bars denote the least significant p-value.

Using a linear mixed effects model, we then compared the effects of conditions on muscle activity during the EVR epoch to identify any impact of the target jump and/or instruction. The model included a four-way interaction between instruction, location of the initial target, target jump, and muscle, with random intercepts for each participant to account for individual variability. Pairwise comparisons were performed on the estimated marginal means to identify significant differences between conditions.

Two statistical patterns emerge from the analysis of mean EMG activity during the EVR epoch (80-120ms; fig. 8B). Firstly, there is a consistently significant difference in EMG between no jump and jump conditions. When arm adduction is required (T1 to T2), adductors exhibit significantly higher activity during jump conditions (i.e. ‘Follow,’ sternal pec. maj.: z = 10.28, p < .001) and abductors exhibit significantly lower activity (i.e. ‘Follow,’ triceps: all z = −3.33, p < .05). The inverse is true when arm abduction is required (T2 to T1). This occurs across all instruction types to varying degrees, even in ‘Ignore’ blocks which necessitate no change in trajectory. Secondly, instruction has an impact on the magnitude of the EVR during jump trials, with participants exhibiting the most prominent EVR during ‘Follow’ jump trials compared to both other blocks (i.e. ‘Follow’ vs. ‘Ignore’, T1 to T2, sternal pec. maj.: z = 3.97, p < .001).

### Stronger EVRs precede earlier online corrections

To investigate the relationship between the EVR and F-RT, we then calculated linear mixed models on an individual muscle basis to investigate the effects of EVR magnitude, conditions, and instruction on F-RT, with random intercepts for each participant to account for individual variability. We found that participants exhibited significantly faster F-RT in response to the target jump during ‘Follow’ blocks than during the other two tasks (all z < −7.77, p < .001; fig. 9A). This pattern is likely due to the EVR magnitude being significantly larger in the ‘Follow’ block followed by the ‘Stop’ and ‘Ignore’ blocks (fig. 7). Pairwise comparisons were performed on the estimated marginal means to identify significant differences between conditions. In line with the EVR pattern observed relative to no jump conditions (fig. 9B), EVR magnitude was predictive of F-RT. For example, an increase in the EVR in the sternal pectoralis major in the ‘Follow’ and ‘Ignore’ blocks was weakly, but significantly, correlated with lower F-RTs in T1 to T2 jumps (all r^2^ > .029, z < −2.91, p < .001), with the inverse being true for T2 to T1 jumps for all instructions (all r^2^ > .032, z > 2.31, p < .05), as is suggested by the direction of the EVR as seen in the group EMG analysis (fig. 9B). Although, individually, EVRs in any one muscle accounted for only a maximum 9.5% of variance in F-RT, a multiple regression analysis of the effect of the mean EVR magnitude of all six muscles on F-RT, showed that, on average, EVR magnitude explained 24.7% of the variance in F-RT (fig. 9C). Collectively, these findings indicate that low-latency online corrections are predicated on the EVR and explain why online corrections toward the jumped target are initiated regardless of block instruction.

**Figure 9.**
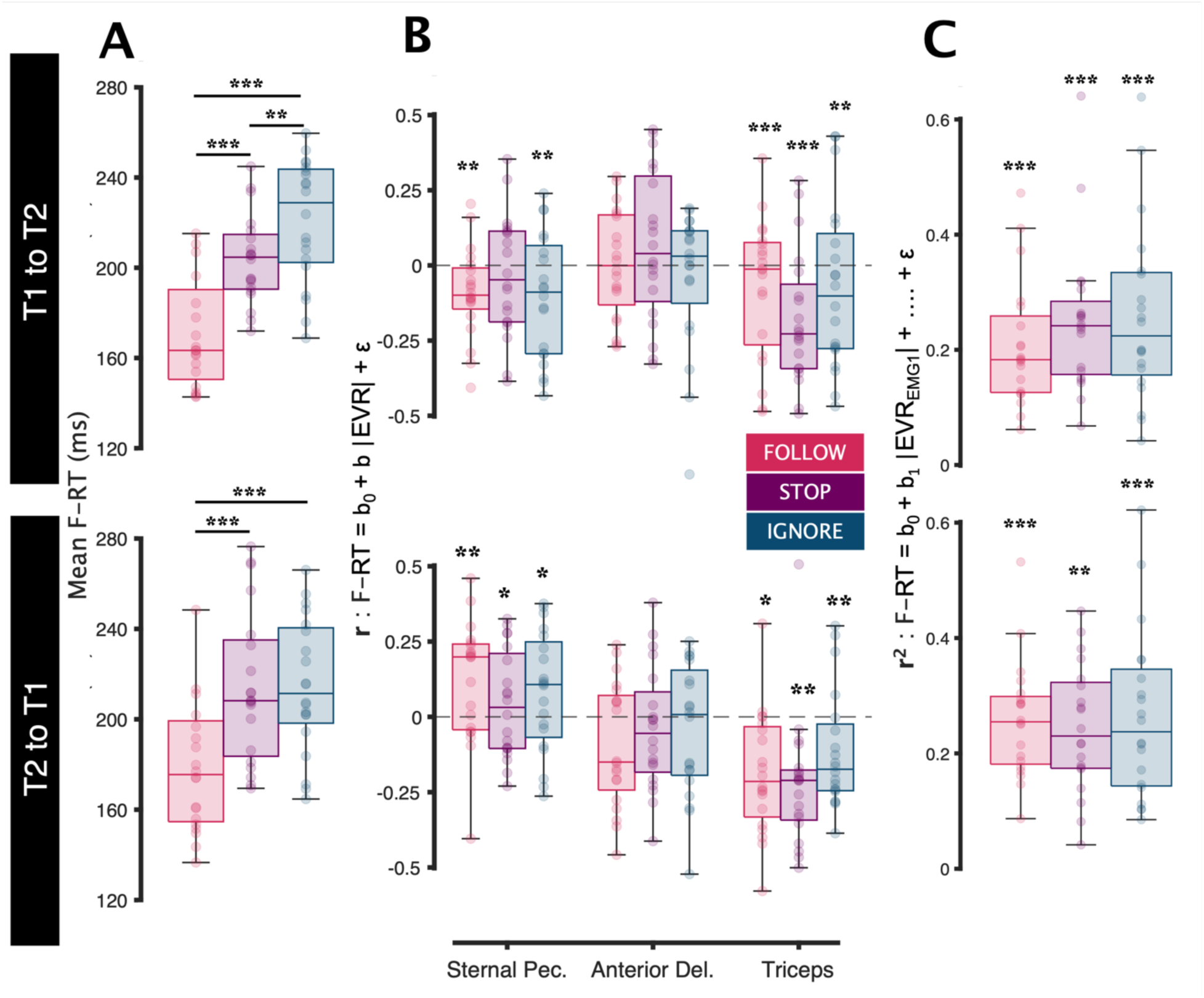
Comparison of F-RT across blocks, and correlation with EVR magnitude: (A) Mean F-RT of jump trials for all participants, excluding trials where F-RT was too early (< 100ms) or did not occur. Asterisks indicate significant findings using a contrast of estimated marginal mean based on the linear mixed effects model used in figure 8. (B) Plotted correlation coefficients between F-RT and mean EMG magnitude during the EVR epoch (80-120ms), reflecting the relationship between EVR magnitude and the direction of movement. Asterisks indicate significant findings using a contrast of estimated marginal mean of the trends based on linear mixed effects models of individual muscles. Degrees of freedom were estimated using the kenward-roger method. The Tukey method was used to adjust p-values. (C) r^2^ for multiple regression analyses of the effect of EVR magnitude of each of the six muscles on F-RT on an individual trial basis for each participant. Asterisks denote significant findings using a Fisher’s combined probability test of the resulting p-values.

### The EVR does not support countermanding

Although we observed that low-latency force changes (ranging from 100 to 150ms after the target jump) typically resulted in kinematic deviation towards the jumped target across all blocks (fig. 6), we also observed that EVR magnitude correlates with F-RT independent of instruction and that EVR magnitude differs between ‘Follow’ blocks and ‘Stop’ blocks (fig. 8). This left open the possibility that the EVR may be contributing to countermanding actions in the ‘Stop’ block. To investigate this, we first compared EMG values across the three instruction types, including both correct and error trials; error trials were included to prevent biasing the analysis, given the disparate error rate between conditions. We found that EMG activity between the ‘Follow’ and ‘Stop’ block did not significantly differ during the EVR epoch in any of the muscles recorded (BF < 3), with the earliest significant difference beginning at a minimum latency of 138ms (fig. 10A), after the EVR.

**Figure 10.**
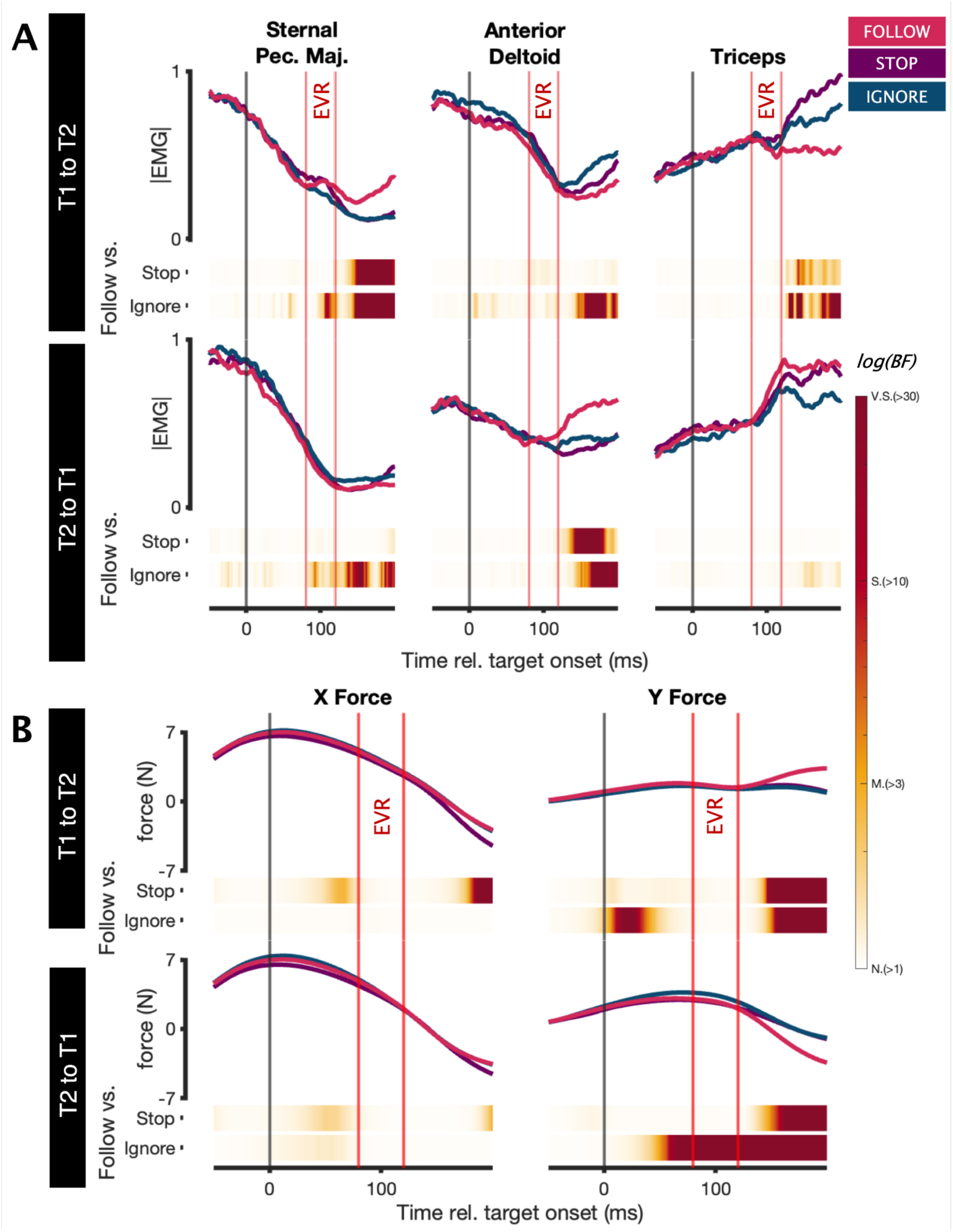
Comparison of EMG activity and force applied to manipulandum across jump conditions in the three different blocks: Comparison of (A) mean group EMG for three muscles and (B) mean force across jump conditions to identify when these measures differ in the three different blocks. BF heatmaps below each graph indicate comparisons between the ‘Follow’ block and each of the other two block types, based on Bayesian two-tailed paired t-test. BF is qualified by the following standard thresholds^30^: N(> 1) = Negligible, M(> 3) = Moderate, S(> 10) = Strong, VS(> 30) = Very Strong. EVR epoch (80-120ms) is denoted by vertical red lines.

As with muscle activity, the initial portion of the force trajectory did not differ between ‘Follow’ and ‘Stop’ blocks, with differences beginning to emerge only ∼150ms following the target jump. Thus, the initial reaction correction beginning ∼110ms after the target jump is relatively similar on ‘Follow’ and ‘Stop’ blocks, moving the hand towards the jumped target. As an aside, this analysis revealed a slight increase in force in the ‘Ignore’ block relative to the ‘Follow’ block during the initial reach (well before the EVR) in one comparison (T2 to T1). This is likely because participants made slightly faster reaches from rest during the ‘Ignore’ block since they were primed to not respond to the target jump and therefore exercised less caution.

## Discussion

Here, we addressed the hypothesis that online corrections to visual targets constitute a distinct class of movements. Such a conceptualization is based on the unique kinematic features of online corrections, including their very short latency and initial stimulus-driven nature. By measuring EMG as well as the hand force on the manipulandum, we find instead support for a nested model of reaching control, composed of distinct phases of subcortical (80-120ms) and cortical (> 130ms) muscle recruitment which occurred at nearly identical latencies during reaching from rest or online correction. The subcortical portion is the EVR, which is time aligned to the stimulus onset rather than movement onset^16^ and spatially aligned to the stimulus; facilitating arm movement towards the new target. However, only during online corrections did this pattern of muscle recruitment lead to low-latency kinematic changes. Thus, an alternative interpretation of previous kinematic results supporting the notion of an ‘automatic pilot’ system for the control of online correction, is that they arose not from earlier muscle recruitment, but rather from task-relevant modulation of these loops and biomechanical considerations related to how quickly consequent forces overcome the arm’s inertia.

### Re-evaluating the model of online corrections

A core prediction of the ‘automatic pilot’ hypothesis is that muscle activity should change sooner after target displacement for online corrections compared to reaches from rest. For example, Smeets, Oostwoud, and Brenner^1^ suggested that online corrections are earlier because there is no need to detect the target de novo following its displacement [but see Brenner and Smeets^34^ for different results from a rapid pointing task]. Conversely, Cluff and Scott^35^ suggested that online corrections are expedited due to the earlier expression of postural adjustments that must precede movements from rest.

When we analyzed movement onset using conventional kinematic measures like acceleration using a standard threshold analysis, we did indeed find earlier onsets for online corrections than movements from rest (Fig. 2). However, a close inspection of manipulandum force and trial-by-trial changes in muscle recruitment revealed essentially identical latencies but larger magnitudes for online corrections than for movements from rest (Figs. 2, 3). This discrepancy highlights the importance of limb biomechanics, and particularly an appreciation of how quickly the forces that arise from changes in muscle recruitment overcome the limb’s inertia. Critically, the impetus^9,36^ and criterion for categorizing online corrections as a distinct class of movement in the previous literature is that they are ‘quick’^2^, ‘fast’^37,38^, or ‘rapid’^39^. Our EMG and force findings are inconsistent with this conception and, by extension, the idea of an ‘automatic pilot’ which functions exclusively in the context of an ongoing movement^3,23,24,38,40^.

Why then was the EVR not observed during previous studies of online corrections, despite occasional use of EMG? Certainly task features appear critical, as noted above, and it is well documented that the EVR^16,41^ and other subcortical reflexes such as express saccades^42^ or responses to other visual perturbations^43,44^ are scaled with task demands and more prevalent during tasks with a degree of urgency. Further, the study of EVRs is predicated on trial-by-trial representations of EMG activity that permit identification of a distinctive band of muscle activity from 80-120ms^15,16,31^; the averaging of EMG activity across multiple trials regardless of reaction time can obscure the EVR and compromise the conclusions that can be drawn. This issue is particularly problematic during online corrections, as in the ‘Follow’ block, as any EVRs would evolve on top of a pre-existing dynamic of muscle recruitment related to the initial reaching movement^31^, and because of the potential for ‘voluntary’ or even anticipatory recruitment to merge into the EVR interval for short-latency trials. However, the EVR is easily dissociated from the subsequent phases of recruitment in the ‘Ignore’ and ‘Stop’ blocks (fig. 7 & 8).

### The EVR underlies the stimulus driven nature of online corrections to visual targets

We investigated the mechanisms underlying the almost irresistible urge to follow a suddenly displaced target in a reaching task. Our results demonstrate that these short-latency online corrections (< 120ms) are preceded by a distinctive phase of muscle recruitment, the express visuomotor response (EVR), which is time locked to the target jump. Here, we show that the magnitude of the EVR in an online correction predicted the timing and magnitude of forces that direct the arm to the target. Further, while the EVR could be muted in magnitude, it persisted even when counter to task goals in the ‘Stop’ and ‘Ignore’ blocks. Thus, both the initial phase of online correction^8,18^ and the EVR (fig. 7 & 8) are automatic and only somewhat muted by task demands. Automatic processing of the jumped target in the ‘Stop’ block emphasizes that errors in this block were more related to irrepressible processing of the jumped target rather than to the jumped target being ignored.

Despite the automaticity of the EVR, the EVR was more pronounced during the ‘Stop’ blocks compared to the ‘Ignore’ block, leading to larger erroneous deviations (fig. 5B) and a shorter latency for those deviations (fig. 9). Presumably, this is because the jumped target still carries behavioral relevance in the ‘Stop’ block, but not in the ‘Ignore’ block; in other words EVRs to task-irrelevant stimuli are more muted, although not completely suppressed^8,45^. This result complements findings that both spatial^32,46,47^ and temporal^48,49^ factors can impact EVR magnitude, and EVRs persistently direct the arm along a path toward a salient visual stimulus in tasks with incongruous stimulus-response mappings^33,47^. Thus, the EVR develops task-relevant forces capable of deviating the arm toward the jumped stimulus in our task, doing so in an automatic fashion.

### A common nested system for visually-guided reaches

Our findings of equivalent timing for reaches whether initiated from rest or during an ongoing movement resonate with models of a nested system for the control of reaching, made up of subcortical and cortical pathways contributing distinct phases of muscle recruitment^32^. In this model the first phase of recruitment, the EVR, is mediated through the superior colliculus (SC) via the tecto-reticulo-spinal pathway. This subcortical pathway has the advantage of being extremely short-latency but the disadvantage of being inexorably stimulus-driven. In monkeys, EVR latency, which is shorter relative to humans^50,51^, is correlated with and preceded by visual bursts in the movement-related layers of the SC^52^. Certain stimuli, including high-contrast^48,53,54^, low-spatial frequency^31,55,56^, and evolutionarily relevant images, like faces^57–59^, preferentially evoke the EVR, activate the human SC, and robustly activate monkey SC neurons within < 50ms. Our observations here add to the evidence that the EVR, and by extension the initial phase of low-latency online corrections to visual targets, are mediated through the tecto-reticulo-spinal pathway. While the nested architecture we propose may well be recruited to targets of other modalities, we speculate that the strong visual response in the SC engenders the stimulus-directed nature to “express” responses which is not seen for auditory^60^ or tactile^61^ modalities.

The second voluntary phase of muscle recruitment, which lags the EVR, is mediated via the canonical cortical motor pathway which, although being slightly slower, has the benefit of being fully flexible to permit any stimulus-response association. This includes the PPC which, through transcranial magnetic stimulation (TMS) and lesions studies, has been classically regarded as the most likely candidate for the ‘automatic pilot’^2,37–39^. Although TMS of human PPC disrupts both online reaching^4,62–65^ and grasping^66–69^ [except when TMS is applied > 100ms after target jump^70,71^], in studies where reaction time was measured, no change was found^4,63^. Instead, TMS of the PPC increased total movement time. The lack of an effect on reaction time suggests that the initial phase of online corrections may have been preserved, possibly because PPC stimulation disrupted the second phase of recruitment without influencing the EVR. It is during this latter interval where muscle recruitment starts to differ if the task demands an incongruent stimulus-response mapping, as an anti-reach paradigm^33^, or in the ‘Stop’ block studied here. In the ‘Stop’ block, the earliest change in muscle recruitment related to actively braking the reach movement occurred ∼20-30ms after the EVR, or ∼140-150ms after the jumped target (fig. 10A). This later latency resembles the timing of an active braking signal on limb muscles when reaching movements are abruptly canceled in standard stop-signal tasks^10^. Thus, when viewed from the perspective of the timing and the inexorable stimulus-driven nature of the EVR, the earliest circuit recruited by a peripheral stimulus can support movement only *toward* a stimulus, and cannot be arbitrarily mapped onto movement cancellation. This perspective differs from a recent report that had reported similar timing for movement initiation and cancellation^13,14^, perhaps because experimental conditions in this study did not favor production of an EVR to the imperative stimulus.

Individuals with optic ataxia following PPC damage exhibit deficits during both online corrections^3,72–74^ and reaching from rest (for review, see Andersen et. al^75^ & Goodale and Milner^76^), particularly to peripheral targets but also, to a lesser degree, central targets. Can these results be reconciled with the notion of nested pathways? As stated in Pisella and colleagues’ ^3^ seminal study on the topic, in their experimental design they were compelled to increase the distance between the initial target (which appeared centrally) and the jumped target because of the patients’ high pointing variability from rest. Notably, none of these studies recorded muscle activity; hence the effect of PPC damage muscle recruitment is unknown. Muscle recordings in patients with optic ataxia, in conjunction with tasks promoting response urgency, are particularly important given recent work in Parkinson’s Disease attributing a preservation of eye and hand movements in interceptive tasks to signaling through the superior colliculus^77^. This work is consistent with other emerging work showing that the pathophysiology of Parkinson’s disease spares EVRs but selectively impacts muscle recruitment during the voluntary interval^78,79^. Whether such selective sparing of EVRs also happens following damage to the PPC and in the context of optic ataxia remains to be determined.

The importance of movement biomechanics and the notion of nested pathways, helps resolve a critical problem in the ‘automatic pilot’ hypothesis pointed out by Sarlegna and Mutha ^39^. The authors questioned why a continuously updated ‘automatic pilot’ capable of implementing an anti-reach from rest would not be able to do so during an ongoing movement; recall that the initial phase of anti-reaches during online corrections is inevitably drawn toward the jumped stimulus^7^. This question is prescient, since it suggests two different modules; one capable and the other incapable of facilitating anti-reaching, with the latter only being active during ongoing movements. However, the EVR is directed toward the stimulus on anti-reach trials from rest^33^, and toward the jumped stimulus even in the ‘Stop’ block. What appears differ is the consequences of this EVR on movement kinematics. During an ongoing movement, the EVR leads to sizable kinematic deviations. From rest, however, the forces arising from the EVR apparently are not sufficient to overcome the arm’s inertia as quickly, even though the EVR starts at exactly the same time. This may be because of a combination of factors, including the EVR being larger in magnitude during ongoing movements and/or distributed to more muscles (fig. 3), because of the force-velocity relationship on actively lengthening muscles^9^, or because the various ligaments and tendons are already tight when the EVR arrives during an ongoing movement. Regardless, for the conditions we studied, we conclude that the timing of the earliest phase of muscle recruitment, on a trial-by-trial basis, is equivalent for movements started from rest and online corrections. This observation reinforces the importance of considering movement biomechanics when trying to infer the reasons why kinematic changes to a movement’s trajectory differ for ongoing movements versus movements from rest. Further, the variability of EVR magnitude, when combined with movement biomechanics, may help explain the progressive increase in response robustness for target displacements introduced later in an ongoing movement^80^. Considerations of task demands are also essential, as EVRs can be generated either well before^81^ or even without subsequent voluntary muscle recruitment^82^. Conversely, robust voluntary recruitment can precede a muted (as seen in the ‘Ignore’ block) or entirely absent EVR, as in non-urgent or delayed response tasks that favour movement accuracy over speed^16^.

### Limitations

Our results do not support the conception of online corrections to visual targets as a distinct class of movements mediated by a rapid ‘automatic pilot’ system. Our main evidence is the equivalent timing of trial-by-trial EMG recruitment and onset of mean manipulandum force toward an imperative stimulus regardless of whether reaching movements are initiated from rest or adjusted in mid-flight, which we argue support a nested system wherein distinct subcortical and cortical loops can be engaged during visually-guided reaching. One clear limitation of our experiments is the absence of causal manipulations. Future studies could employ neurostimulation techniques like transcranial magnetic stimulation to test the architecture of the nested system. Further, subjects knew that relevant visual stimuli would be presented at only two fairly adjacent spatial locations; future extensions of this work should explore workspaces with a higher degree of target uncertainty. Finally, we did not measure eye movements, which could theoretically reveal relationships between express saccades and the EVR. Regardless, our results emphasize the importance of recording muscle activity and/or force measures in addition to movement kinematics to ensure accurate interpretation of experimental data.

## Methods

### Participants

All procedures were approved by the Health and Science Research Ethics Board at Western University, London, Ontario, Canada, and conformed to standards set by the Declaration of Helsinki. Twenty university students (female: 15, male: 5; mean age: 19.15 ± 2.08; right-handed: 19, left-handed: 1) were recruited using the Psychology Research Participant Pool, with compensation provided as credits required for course completion. All provided written consent, reported normal or corrected to normal vision and no neurological disorders, completed the experiment, and are included in the analysis.

### Apparatus

Participants made visually-guided reaching movements in a Kinarm Endpoint Robot (fig. 1A; BKIN technologies, Kingston, Ontario, Canada) integrated with a high-performance Propixx projector (VPixx, Saint-Bruno, Quebec, Canada) that ensured reliable event timing. A photodiode provided additional verification of stimulus timing. Behavioural tasks were generated using Stateflow and Simulink within MATLAB (version R2015a, MathWorks Inc., Natick, Massachusetts, United States of America).

All participants, regardless of handedness, completed the task with their right arm using a manipulandum under a horizontal display surface at approximately shoulder level when seated. The display occluded the participants’ hand, but did not impact their ability to perform online reach corrections ^40,83^. Surface electromyography (EMG; Delsys Inc. Bagnoli-8 system, Boston, Massachusetts, United States of America) recordings of the following six right arm muscles, identified through palpation, were acquired: clavicular and sternal heads of pectoralis major, anterior and posterior deltoids, short head of biceps, and medial head of triceps (fig. 1B). EMG data was filtered using a high and low pass filter of 20 and 450Hz. All data was recorded at 1000Hz by the Kinarm computer.

### Behavioural task

Participants initiated trials by aligning their cursor (r = 1cm), controlled by the Kinarm manipulandum, within the startpoint (r = 1cm). Following a hold of 750ms, the startpoint disappeared. Following a randomised time of 250ms + 1:1000ms (total hold = 1000:2000ms), a target (r = 3cm) appeared randomly in either the T1 or T2 location (fig. 1C). These target locations were chosen to be in the preferred direction for EVRs on the clavicular and sternal heads of pectoralis major^46,84^. Temporal uncertainty was used to reduce anticipatory reaches. If the participant’s hand left the startpoint location before target onset, the trial was terminated and repeated. In 66% of trials, no jump occurred and the trial ended when the cursor reached the target (fig. 1D). In 33% of trials, upon movement initiation (centre of cursor > 1cm away from centre of startpoint) the target jumped to the other location (fig. 1E-G).

In different blocks, we used identical visual stimuli but instructed participants to perform different motor adjustments in response to a target jump. By using the same stimuli, we could ensure that observed differences stemmed solely from top-down cognitive task setting rather than bottom-up stimuli related factors. If the target jumped (E) during ‘Follow’ blocks, participants were instructed to ‘Reach to **NEW** target,’ (F) during ‘Stop’ blocks, they were instructed to ‘**STOP**’ the ongoing movement and hold their hand in place, (G) and during ‘Ignore’ blocks, they were instructed to ‘Reach to **OLD** target.’ On jump trials, the trial ended in ‘Follow’ blocks when the cursor touched the new target (fig. 1E), in ‘Stop’ blocks 1500ms following target jump (fig. 1F), and in ‘Ignore’ blocks when the cursor touched the old target location (fig. 1G).

A practice session of 78 trials was completed before the experiment, including all trial types broken over 6 blocks with alternating instructions, followed by a timed 3-minute break. The experiment consisted of 864 trials, including 192 no jump (66%) and 96 jump (33%) trials for each instruction type. The experiment was broken up into 9 blocks of 96 trials, after each of which the instruction was alternated. The order of the block instructions was pseudorandomized across participants, then repeated sequentially over the experiment, resulting in 3 blocks for each instruction type (i.e: Ignore-Follow-Stop-Ignore-Follow-Stop…). A timed 5-minute break followed every 3 blocks of the experiment. The entire experiment took about 90 minutes.

### Data alignment

All preprocessing and analyses, unless otherwise stated, were completed using MATLAB (version R2021a, MathWorks Inc., Natick, Massachusetts, United States of America). On each trial, an unseen white stimulus appeared simultaneously with the first target onset and disappeared once the participant’s cursor left the startpoint (the same time the second target appeared and the first target disappeared on jump trials). The onset and offset of this white stimulus was detected by a diode. Kinematic and EMG data were aligned on a per-trial basis to the diode onset (initial target onset) or offset (target jump), depending on whether we wished to investigate the initial reach or the online reach correction, respectively.

### Analyses of movement kinematics and forces

We evaluated performance to ensure participants completed the task as instructed. Jump trials were denoted as error trials if the participant failed to follow the block instruction. Using coordinates from the Kinarm manipulandum, trials were marked as error in ‘Follow’ blocks if the cursor made contact with the initial target location, in ‘Stop’ blocks if the cursor made contact with either of the target locations, and in ‘Ignore’ blocks if the cursor made contact with the new target (fig. 4A). No trials were excluded on the basis of these performance classifications in any of our analyses.

In addition to recording kinematic measures such as position data and its derivatives, we also analyzed the force exerted by the hand on the manipulandum, which was measured via force transducers. Force data was filtered using a second-order low-pass Butterworth filter of 60Hz. Force and kinematic data was then transformed spatially by rotating the workspace - 157.5° around the centre of the startpoint (see fig. 5A for a graphical depiction of this rotation). This ensured that the x-axis denoted movement between the startpoint and midpoint between the targets, while the perpendicular y-axis denoted deviation of the trajectory towards one of the two targets.

For each participant, we used data from no jump trials to determine the mean force trajectory in the x and y dimensions, as well as the standard deviation of the Euclidean differences relative to this mean. Force reaction time (F-RT) of each jump trial was determined by identifying the time point when the Euclidean difference exceeded one standard deviation, relative to the complementary no jump condition. We calculated F-RTs following target jumps for both jump and no jump trials (as a control) to determine the earliest F-RT onset latency. We found that the earliest discernible difference between the two distributions was ∼100ms. Therefore, for all analyses we excluded trials where F-RT occurred prior to 100ms, excluding on average 14.6% of jump trials in ‘Follow’ blocks, 16.7% in ‘Stop’ blocks, and 14.7% in ‘Ignore’ blocks. Inclusion or exclusion of these trials did not change our result, however, we opted to exclude them as a control of anticipatory reach adjustments. Notably, these anticipatory adjustments were approximately just as likely (∼15%) during jump and no jump trials since conditions were intermixed and randomized.

We did not exclude any trials due to any task-specific factors such as any error in the reach adjustment (reaching to the wrong target or performing the wrong reach adjustment) or a delay or absence of a response. By removing any subjective criteria, we aimed to objectively assess the impact of muscle recruitment on force and kinematics. The exception to this is for the analysis of F-RT, where we excluded trials without a detected F-RT (which happened when the participant’s hand did not deviate from the no-jump trajectory). This happened on average in an additional 1.3% of jump trials in ‘Follow’ blocks, 3.4% in ‘Stop’ blocks, and 24.3% in ‘Ignore’ blocks.

For each participant the mean Euclidean difference of force was then calculated for both no jump and jump trials, relative to the mean no jump trajectory. A group analysis was conducted with a timewise one-tailed paired t-test using the function BayesFactor v.2.3.0 ^85^, to quantify the likelihood of the model that force in jump trials deviated more than no jump trials, relative to the mean no jump force trajectory. The same analysis was used to compare the force of the initial reach with baseline force.

### Analysis of muscle activity

Offline, EMG data was full-wave rectified and then filtered by a 7-point moving average filter. Since participants initiated a target jump during jump trials when their hand left the startpoint, we aligned each trial’s data to the time point for both trials when the target did jump on jump trials, and when it would have jumped on no-jump trials. This allowed us to compare EMG data during the EVR epoch (80-120ms) of jump trials with the same time point during trials where the target did not jump. The relationship between the mean EMG magnitude during the EVR epoch, muscle, condition, and F-RT was analyzed using linear mixed effects models, performed using R (v4.4.0, R Foundation for Statistical Computing, Vienna, Austria). Specific models are discussed in the results section. A Pearson’s correlation was used to visualize the relationship between EMG magnitude on F-RT. Additionally, group analyses were conducted using a timewise two-tailed paired Bayesian t-test, to quantify the likelihood of a difference of these in EMG magnitude across trial types.

## Acknowledgements

This work is supported by Discovery Grants from the Natural Sciences and Engineering Research Council of Canada (NSERC) to BDC (RGPIN 311680 and 04394-2021) and MAG (RGPIN-2017-04088).

## Author contributions

Conceptualization — DYM, MAG, BDC; data curation — DYM; formal analysis — DYM; funding acquisition — MAG, BDC; investigation — DYM; methodology — DYM, MAG, BDC; project administration — MAG, BDC; resources — MAG, BDC; software — DYM; supervision — MAG, BDC; validation — MAG, BDC; visualization — DYM; original draft — DYM; review and editing — DYM, MAG, BDC.

## STAR Methods

### Key resources table

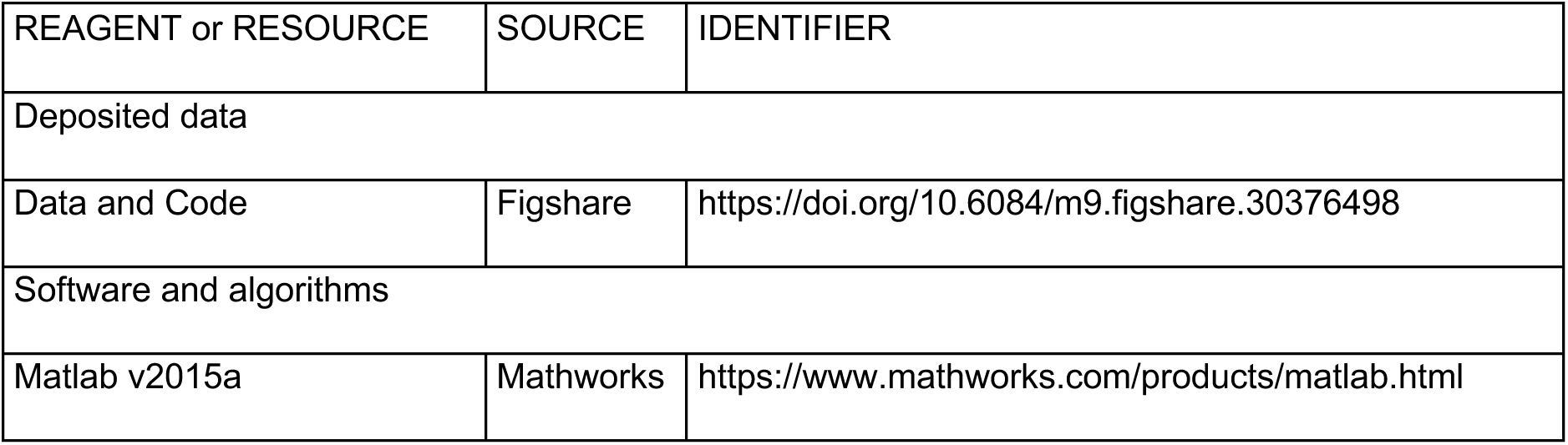

| REAGENT or RESOURCE | SOURCE | IDENTIFIER |
| --- | --- | --- |
| Deposited data |  |  |
| Data and Code | Figshare | https://doi.org/10.6084/m9.figshare.30376498 |
| Software and algorithms |  |  |
| Matlab v2015a | Mathworks | https://www.mathworks.com/products/matlab.html |

## Abbreviations

BF: Bayes factor
EMG: Electromyography
EVR: Express visuomotor response
F-RT: Force reaction time
PPC: Posterior parietal cortex
SC: Superior colliculus
TMS: Transcranial magnetic stimulation

## References

1. Smeets, J.B., Oostwoud Wijdenes, L., and Brenner, E. (2016). Movement Adjustments Have Short Latencies Because There is No Need to Detect Anything. Motor Control 20, 137–148.

2. Gomi, H. (2008). Implicit online corrections of reaching movements. Curr Opin Neurobiol 18, 558–564.

3. Pisella, L., Gréa, H., Tilikete, C., Vighetto, A., Desmurget, M., Rode, G., Boisson, D., and Rossetti, Y. (2000). An “automatic pilot” for the hand in human posterior parietal cortex: toward reinterpreting optic ataxia. Nat. Neurosci. 3, 729–736.

4. Desmurget, M., Epstein, C.M., Turner, R.S., Prablanc, C., Alexander, G.E., and Grafton, S.T. (1999). Role of the posterior parietal cortex in updating reaching movements to a visual target. Nat. Neurosci. 2, 563–567.

5. Day, B.L., and Brown, P. (2001). Evidence for subcortical involvement in the visual control of human reaching. Brain 124, 1832–1840.

6. Reynolds, R.F., and Day, B.L. (2012). Direct visuomotor mapping for fast visually-evoked arm movements. Neuropsychologia 50, 3169–3173.

7. Day, B.L., and Lyon, I.N. (2000). Voluntary modification of automatic arm movements evoked by motion of a visual target. Exp Brain Res 130, 159–168.

8. Striemer, C.L., Yukovsky, J., and Goodale, M.A. (2010). Can intention override the “automatic pilot”? Exp. Brain Res. 202, 623–632.

9. Soechting, J.F., and Lacquaniti, F. (1983). Modification of trajectory of a pointing movement in response to a change in target location. J Neurophysiol 49, 548–564.

10. Atsma, J., Maij, F., Gu, C., Medendorp, W.P., and Corneil, B.D. (2018). Active Braking of Whole-Arm Reaching Movements Provides Single-Trial Neuromuscular Measures of Movement Cancellation. J Neurosci 38, 4367–4382.

11. Raud, L., and Huster, R.J. (2017). The Temporal Dynamics of Response Inhibition and their Modulation by Cognitive Control. Brain Topogr 30, 486–501.

12. Raud, L., Westerhausen, R., Dooley, N., and Huster, R.J. (2020). Differences in unity: The go/no-go and stop signal tasks rely on different mechanisms. Neuroimage 210, 116582.

13. Novembre, G., and Iannetti, G.D. (2021). Towards a unified neural mechanism for reactive adaptive behaviour. Prog. Neurobiol. 204, 102115.

14. Du, Y., Forrence, A.D., Metcalf, D.M., and Haith, A.M. (2024). Action initiation and action inhibition follow the same time course when compared under matched experimental conditions. J Neurophysiol 131, 757–767.

15. Corneil, B.D., Olivier, E., and Munoz, D.P. (2004). Visual responses on neck muscles reveal selective gating that prevents express saccades. Neuron 42, 831–841.

16. Pruszynski, J.A., King, G.L., Boisse, L., Scott, S.H., Flanagan, J.R., and Munoz, D.P. (2010). Stimulus-locked responses on human arm muscles reveal a rapid neural pathway linking visual input to arm motor output. Eur. J. Neurosci. 32, 1049–1057.

17. Contemori, S., Loeb, G.E., Corneil, B.D., Wallis, G., and Carroll, T.J. (2022). Symbolic cues enhance express visuomotor responses in human arm muscles at the motor planning rather than the visuospatial processing stage. J Neurophysiol 128, 494–510.

18. Gritsenko, V., and Kalaska, J.F. (2010). Rapid online correction is selectively suppressed during movement with a visuomotor transformation. J Neurophysiol 104, 3084–3104.

19. Paulignan, Y., Jeannerod, M., MacKenzie, C., and Marteniuk, R. (1991). Selective perturbation of visual input during prehension movements. 2. The effects of changing object size. Exp. Brain Res. 87, 407–420.

20. Paulignan, Y., MacKenzie, C., Marteniuk, R., and Jeannerod, M. (1991). Selective perturbation of visual input during prehension movements. 1. The effects of changing object position. Exp. Brain Res. 83, 502–512.

21. Brenner, E., and Smeets, J.B. (1997). Fast Responses of the Human Hand to Changes in Target Position. J Mot Behav 29, 297–310.

22. Reichenbach, A., Thielscher, A., Peer, A., Bülthoff, H.H., and Bresciani, J.-P. (2009). Seeing the hand while reaching speeds up on-line responses to a sudden change in target position. J Physiol 587, 4605–4616.

23. Brière, J., and Proteau, L. (2011). Automatic movement error detection and correction processes in reaching movements. Exp Brain Res 208, 39–50.

24. Prablanc, C., and Martin, O. (1992). Automatic control during hand reaching at undetected two-dimensional target displacements. J Neurophysiol 67, 455–469.

25. Oostwoud Wijdenes, L., Brenner, E., and Smeets, J.B.J. (2011). Fast and fine-tuned corrections when the target of a hand movement is displaced. Exp Brain Res 214, 453–462.

26. de Brouwer, A.J., and Spering, M. (2022). Eye-hand coordination during online reach corrections is task dependent. J Neurophysiol 127, 885–895.

27. Oostwoud Wijdenes, L., Brenner, E., and Smeets, J.B.J. (2014). Analysis of methods to determine the latency of online movement adjustments. Behav Res Methods 46, 131–139.

28. Brenner, E., and Smeets, J.B.J. (2019). How Can You Best Measure Reaction Times? J Mot Behav 51, 486–495.

29. Dienes, Z. (2011). Bayesian Versus Orthodox Statistics: Which Side Are You On? Perspect Psychol Sci 6, 274–290.

30. Jeffreys, H. (1998). The Theory of Probability (OUP Oxford).

31. Kozak, R.A., Kreyenmeier, P., Gu, C., Johnston, K., and Corneil, B.D. (2019). Stimulus-Locked Responses on Human Upper Limb Muscles and Corrective Reaches Are Preferentially Evoked by Low Spatial Frequencies. eNeuro 6. 10.1523/ENEURO.0301-19.2019.

32. Contemori, S., Loeb, G.E., Corneil, B.D., Wallis, G., and Carroll, T.J. (2023). Express Visuomotor Responses Reflect Knowledge of Both Target Locations and Contextual Rules during Reaches of Different Amplitudes. J. Neurosci. 43, 7041–7055.

33. Gu, C., Wood, D.K., Gribble, P.L., and Corneil, B.D. (2016). A Trial-by-Trial Window into Sensorimotor Transformations in the Human Motor Periphery. J. Neurosci. 36, 8273–8282.

34. Brenner, E., and Smeets, J.B.J. (2022). Having several options does not increase the time it takes to make a movement to an adequate end point. Exp Brain Res 240, 1849–1871.

35. Cluff, T., and Scott, S.H. (2016). Online Corrections are Faster Because Movement Initiation Must Disengage Postural Control. Motor Control 20, 162–170.

36. Carlton, L.G. (1981). Processing visual feedback information for movement control. J Exp Psychol Hum Percept Perform 7, 1019–1030.

37. Archambault, P.S., Ferrari-Toniolo, S., Caminiti, R., and Battaglia-Mayer, A. (2015). Visually-guided correction of hand reaching movements: The neurophysiological bases in the cerebral cortex. Vision Res 110, 244–256.

38. Gaveau, V., Pisella, L., Priot, A.-E., Fukui, T., Rossetti, Y.R., Pélisson, D., and Prablanc, C. (2014). Automatic online control of motor adjustments in reaching and grasping. Neuropsychologia 55, 25–40.

39. Sarlegna, F.R., and Mutha, P.K. (2015). The influence of visual target information on the online control of movements. Vision Res 110, 144–154.

40. Gosselin-Kessiby, N., Messier, J., and Kalaska, J.F. (2008). Evidence for automatic on-line adjustments of hand orientation during natural reaching movements to stationary targets. J Neurophysiol 99, 1653–1671.

41. Weerdesteyn, V., Kearsley, S.L., Cecala, A.L., Macpherson, E.A., and Corneil, B.D. (2024). Startling acoustic stimuli hasten reflexive choice reaching tasks by strengthening, but not changing the timing of, express visuomotor responses. bioRxiv, 2024.07.01.601510. 10.1101/2024.07.01.601510.

42. Oor, E.E., Stanford, T.R., and Salinas, E. (2023). Stimulus salience conflicts and colludes with endogenous goals during urgent choices. iScience 26, 106253.

43. Nashed, J.Y., Crevecoeur, F., and Scott, S.H. (2012). Influence of the behavioral goal and environmental obstacles on rapid feedback responses. J Neurophysiol 108, 999–1009.

44. Franklin, D.W., and Wolpert, D.M. (2008). Specificity of reflex adaptation for task-relevant variability. J Neurosci 28, 14165–14175.

45. Brenner, E., de Jonge, L.M., Tiems, N.J.J., Rosenquist, G., Wiggers, T., Smeets, J.B.J., and Crowe, E.M. (2025). Can you learn not to respond to irrelevant motion while making fast arm movements? PLoS One 20, e0332171.

46. Selen, L.P.J., Corneil, B.D., and Medendorp, W.P. (2023). Single-Trial Dynamics of Competing Reach Plans in the Human Motor Periphery. J Neurosci 43, 2782–2793.

47. Gu, C., Pruszynski, J.A., Gribble, P.L., and Corneil, B.D. (2018). Done in 100 ms: path-dependent visuomotor transformation in the human upper limb. J Neurophysiol 119, 1319–1328.

48. Kozak, R.A., and Corneil, B.D. (2021). High-contrast, moving targets in an emerging target paradigm promote fast visuomotor responses during visually guided reaching. J. Neurophysiol. 126, 68–81.

49. Contemori, S., Loeb, G.E., Corneil, B.D., Wallis, G., and Carroll, T.J. (2021). Trial-by-trial modulation of express visuomotor responses induced by symbolic or barely detectable cues. J Neurophysiol 126, 1507–1523.

50. Cecala, A.L., Kozak, R.A., Pruszynski, J.A., and Corneil, B.D. (2023). Done in 65 ms: Express Visuomotor Responses in Upper Limb Muscles in Rhesus Macaques. eNeuro 10. 10.1523/ENEURO.0078-23.2023.

51. Goonetilleke, S.C., Katz, L., Wood, D.K., Gu, C., Huk, A.C., and Corneil, B.D. (2015). Cross-species comparison of anticipatory and stimulus-driven neck muscle activity well before saccadic gaze shifts in humans and nonhuman primates. J Neurophysiol 114, 902–913.

52. Rezvani, S., and Corneil, B.D. (2008). Recruitment of a head-turning synergy by low-frequency activity in the primate superior colliculus. J. Neurophysiol. 100, 397–411.

53. Chen, C.-Y., and Hafed, Z.M. (2018). Orientation and Contrast Tuning Properties and Temporal Flicker Fusion Characteristics of Primate Superior Colliculus Neurons. Front. Neural Circuits 12, 58.

54. Schneider, K.A., and Kastner, S. (2005). Visual responses of the human superior colliculus: a high-resolution functional magnetic resonance imaging study. J. Neurophysiol. 94, 2491–2503.

55. Chen, C.-Y., Sonnenberg, L., Weller, S., Witschel, T., and Hafed, Z.M. (2018). Spatial frequency sensitivity in macaque midbrain. Nat. Commun. 9, 2852.

56. Zhang, P., Zhou, H., Wen, W., and He, S. (2015). Layer-specific response properties of the human lateral geniculate nucleus and superior colliculus. Neuroimage 111, 159–166.

57. Wang, Y.C., Bianciardi, M., Chanes, L., and Satpute, A.B. (2020). Ultra High Field fMRI of Human Superior Colliculi Activity during Affective Visual Processing. Sci. Rep. 10, 1331.

58. Mekhaiel, D.Y., Goodale, M.A., and Corneil, B.D. (2024). Rapid integration of face detection and task set in visually guided reaching. Eur J Neurosci 60, 5328–5347.

59. Yu, G., Katz, L.N., Quaia, C., Messinger, A., and Krauzlis, R.J. (2024). Short-latency preference for faces in primate superior colliculus depends on visual cortex. Neuron 112, 2814–2822.e4.

60. Shafiq, R., Stuart, G.W., Sandbach, J., Maruff, P., and Currie, J. (1998). The gap effect and express saccades in the auditory modality. Exp Brain Res 118, 221–229.

61. Reschechtko, S., and Pruszynski, J.A. (2020). Voluntary modification of rapid tactile-motor responses during reaching differs from its visuomotor counterpart. J Neurophysiol 124, 284–294.

62. Johnson, H., and Haggard, P. (2005). Motor awareness without perceptual awareness. Neuropsychologia 43, 227–237.

63. Reichenbach, A., Bresciani, J.-P., Peer, A., Bülthoff, H.H., and Thielscher, A. (2011). Contributions of the PPC to online control of visually guided reaching movements assessed with fMRI-guided TMS. Cereb. Cortex 21, 1602–1612.

64. Breveglieri, R., Borgomaneri, S., Filippini, M., Tessari, A., Galletti, C., Davare, M., and Fattori, P. (2023). Complementary contribution of the medial and lateral human parietal cortex to grasping: a repetitive TMS study. Cereb. Cortex 33, 5122–5134.

65. Della-Maggiore, V., Malfait, N., Ostry, D.J., and Paus, T. (2004). Stimulation of the posterior parietal cortex interferes with arm trajectory adjustments during the learning of new dynamics. J. Neurosci. 24, 9971–9976.

66. Glover, S., Miall, R.C., and Rushworth, M.F.S. (2005). Parietal rTMS disrupts the initiation but not the execution of on-line adjustments to a perturbation of object size. J. Cogn. Neurosci. 17, 124–136.

67. Tunik, E., Frey, S.H., and Grafton, S.T. (2005). Virtual lesions of the anterior intraparietal area disrupt goal-dependent on-line adjustments of grasp. Nat. Neurosci. 8, 505–511.

68. Rice, N.J., Tunik, E., Cross, E.S., and Grafton, S.T. (2007). On-line grasp control is mediated by the contralateral hemisphere. Brain Res. 1175, 76–84.

69. Rice, N.J., Tunik, E., and Grafton, S.T. (2006). The anterior intraparietal sulcus mediates grasp execution, independent of requirement to update: new insights from transcranial magnetic stimulation. J. Neurosci. 26, 8176–8182.

70. Savoie, F.-A., Dallaire-Jean, L., Thénault, F., Whittingstall, K., and Bernier, P.-M. (2020). Single-Pulse TMS over the Parietal Cortex Does Not Impair Sensorimotor Perturbation-Induced Changes in Motor Commands. eNeuro 7. 10.1523/ENEURO.0209-19.2020.

71. Marigold, D.S., Lajoie, K., and Heed, T. (2019). No effect of triple-pulse TMS medial to intraparietal sulcus on online correction for target perturbations during goal-directed hand and foot reaches. PLoS One 14, e0223986.

72. Gréa, H., Pisella, L., Rossetti, Y., Desmurget, M., Tilikete, C., Grafton, S., Prablanc, C., and Vighetto, A. (2002). A lesion of the posterior parietal cortex disrupts on-line adjustments during aiming movements. Neuropsychologia 40, 2471–2480.

73. Rossit, S., McIntosh, R.D., Malhotra, P., Butler, S.H., Muir, K., and Harvey, M. (2012). Attention in action: evidence from on-line corrections in left visual neglect. Neuropsychologia 50, 1124–1135.

74. Schaefer, S.Y., Mutha, P.K., Haaland, K.Y., and Sainburg, R.L. (2012). Hemispheric specialization for movement control produces dissociable differences in online corrections after stroke. Cereb Cortex 22, 1407–1419.

75. Andersen, R.A., Andersen, K.N., Hwang, E.J., and Hauschild, M. (2014). Optic ataxia: from Balint’s syndrome to the parietal reach region. Neuron 81, 967–983.

76. Milner, A.D., and Goodale, M.A. (2008). Two visual systems re-viewed. Neuropsychologia 46, 774–785.

77. Fooken, J., Patel, P., Jones, C.B., McKeown, M.J., and Spering, M. (2022). Preservation of Eye Movements in Parkinson’s Disease Is Stimulus- and Task-Specific. J Neurosci 42, 487–499.

78. Billen, L.S., Nonnekes, J., Corneil, B.D., and Weerdesteyn, V. (2024). Lower-limb express visuomotor responses are spared in Parkinson’s Disease during step initiation from a stable position. bioRxiv, 2024.11.29.625631. 10.1101/2024.11.29.625631.

79. Gilchrist, M., Kozak, R.A., Prenger, M., Anello, M., Van Hedger, K., MacDonald, P.A., and Corneil, B.D. (2024). Parkinson’s Disease affects the contextual control, but not the expression, of a rapid visuomotor response that initiates visually-guided reaching: Evidence for multiple, interacting motor pathways and implications for motor symptoms in Parkinson’s Disease. bioRxiv, 2024.11.27.625399. 10.1101/2024.11.27.625399.

80. Gritsenko, V., Yakovenko, S., and Kalaska, J.F. (2009). Integration of predictive feedforward and sensory feedback signals for online control of visually guided movement. J Neurophysiol 102, 914–930.

81. Wood, D.K., Gu, C., Corneil, B.D., Gribble, P.L., and Goodale, M.A. (2015). Transient visual responses reset the phase of low-frequency oscillations in the skeletomotor periphery. Eur J Neurosci 42, 1919–1932.

82. Kearsley, S.L., Cecala, A.L., Kozak, R.A., and Corneil, B.D. (2022). Express arm responses appear bilaterally on upper-limb muscles in an arm choice reaching task. J Neurophysiol 127, 969–983.

83. Goodale, M.A., Pelisson, D., and Prablanc, C. (1986). Large adjustments in visually guided reaching do not depend on vision of the hand or perception of target displacement. Nature 320, 748–750.

84. Gu, C., Pruszynski, J.A., Gribble, P.L., and Corneil, B.D. (2019). A rapid visuomotor response on the human upper limb is selectively influenced by implicit motor learning. J Neurophysiol 121, 85–95.

85. Krekelberg, B. (2022). BayesFactor: Release 2022 (v2.3.0) (Zenodo) 10.5281/ZENODO.7006300.

